# Task-Based Functional Connectomes Predict Cognitive Phenotypes Across Psychiatric Disease

**DOI:** 10.1101/638825

**Authors:** Daniel S. Barron, Siyuan Gao, Javid Dadashkarimi, Abigail S. Greene, Marisa N. Spann, Stephanie Noble, Evelyn Lake, John Krystal, R. Todd Constable, Dustin Scheinost

## Abstract

**Importance:** We show that three common approaches to clinical deficits (cognitive phenotype, disease group, disease severity) each offer useful and perhaps complimentary explanations for the brain’s underlying functional architecture as affected by psychiatric disease.

**Objective:** To understand how different clinical frameworks are represented in the brain’s functional connectome.

**Design:** We use an openly available dataset to create predictive models based on multiple connectomes built from task-based functional MRI data. We use these models to predict individual traits corresponding to multiple cognitive constructs across disease category. We also show that these same connectomes statistically differ depending on disease category and symptom burden.

**Setting:** This was a population-based study with data collected in UCLA.

**Participants:** Healthy adults were recruited by community advertisements from the Los Angeles area. Participants with adult ADHD, bipolar disorder, and schizophrenia were recruited using a patient-oriented strategy involving outreach to local clinics and online portals (separate from the methods used to recruit healthy volunteers)

## INTRODUCTION

Each patient is uniquely complex. Effective clinical care requires a framework that reduces this complexity to trends that a clinician can reliably identify and pair with treatments. Depending on their clinical framework, a clinician might assess specific facets of a patient’s cognitive phenotype like working memory or executive function; might ask about a pattern of symptoms and try to diagnose a patient within a disease group; or might assess a patient’s overall disease severity and decide whether they require inpatient treatment. Each of these three frameworks— phenotype, disease group, and disease severity—can help guide a clinical evaluation, but which is most appropriate to understand a patient’s deficit?

Clinicians have long used symptom-based groups in diagnosis and treatment, however the rise of pharmacotherapy and neurobiology outstripped the precision of symptoms, which can have diverse etiologies (Barron, 2019; Barron et al., 2018). Group-level brain imaging studies have shown that the same neural centers (Goodkind et al., 2015; Vanasse et al., 2018) are implicated in a range of psychiatric diseases, recapitulating genetic studies that undermine the biologic validity of symptom-based groupings. Disease severity is also symptom-based and represents a clinical endpoints in treatment trials.

Cognitive phenotypes that probe discrete neural systems commonly implicated in disease states promise greater validity (Insel, Sahakian, Voon, Nye, & Brown, 2012; Insel et al., 2010). This strategy assumes that clinically-relevant deficits can be identified as a deviation from a central, healthy tendency. Concepts of “health” and “disease” have begun to be seen as statistical consequences of group-level studies, but not reflective of biology. A wide range of brain imaging, behavioral, and genomic studies have shown pervasive phenotypic variability and overlapping distributions at the population level (Cross-Disorder Group of the Psychiatric Genomics Consortium, 2013; Holmes & Patrick, 2018). The clinical question, then, is how to best account for individual and population-level variability and define “deficits” in such a way that they can be identified and alleviated?

We choose to evaluate clinically relevant phenotypes. We performed a series of predictive and explanatory analyses of the brain’s functional connectome (Shen et al., 2017). Functional connectomes have been shown to be unique to an individual (Finn et al., 2015), stable over a lifespan (Horien, Shen, Scheinost, & Constable, 2019), and predictive of clinical and cognitive traits in novel subjects (Dubois, Galdi, Paul, & Adolphs, 2018; Rosenberg et al., 2016). Though connectomes are typically based on resting-state fMRI, task-based fMRI has been shown to improve the prediction of individual cognitive traits and more clearly delineate brain-behavior relationships (Greene, Gao, Scheinost, & Constable, 2018).

To this end, we use connectome-based predictive modeling (CPM) to identify patterns of functional connectivity during a series of fMRI tasks that predict individual differences in cognitive phenotype. We show that models based on these networks involve edges from all over the brain, generalize to previously unseen individuals independent of diagnosis, and define overlapping phenotypic distributions. Notwithstanding these results, a mass multivariate analysis of the same task-based functional connectomes demonstrates differences in disease group and disease severity. Together, these analyses demonstrate that the brain’s functional architecture does not reorganize with mental illness, but that variance within the similar spatially defined, highly complex brain networks account for cognitive phenotype, disease group, and disease severity.

We conclude that three common frameworks for characterizing clinical deficits (cognitive phenotype, disease group, and disease severity) offer useful and perhaps complimentary explanations for the brain’s underlying functional architecture affected by psychiatric disease.

## METHODS

The overarching goal was to evaluate whether the brain’s task-based functional connectivity could predict performance on cognitive constructs across diagnostic categories. To this end, we applied the latest connectome-based predictive modeling (CPM) algorithms to a transdiagnostic dataset gathered by the UCLA Consortium for Neuropsychiatric Phenomics (CNP).

### CNP data set

The CNP dataset (Poldrack et al., 2016) is available through the OpenfMRI project (http://openfmri.org) (Gorgolewski et al., 2016) and includes structural, resting-state functional, and task-based functional neuroimaging (M.R.I.); extensive neuropsychologic assessments and neurocognitive tasks; and demographic information including biologic sex, age, education, and medication history from healthy controls (n=130) and patients with schizophrenia (n=50), bipolar disorder (n=49), and ADHD (n=43; diagnosis by reference to the Structured Clinical Interview for DSM-IV(First, Spitzer, Gibbon, & Williams, 2002)). This dataset has been described in detail elsewhere (Poldrack et al., 2016).

### CNP Participant Selection

We restricted this larger sample to subjects who had full-brain structural images and task-based functional MRI acquisitions during the balloon analog risk task (BART), Paired Associative Memory encoding (PAM-E), Paired Associative Memory retrieval (PAM-R), Spatial Working Memory Capacity (SCAP), Stop Signal (SS), and Task Switching (TS). We excluded 97 subjects (54 controls, 18 schizophrenia, 14 bipolar, and 11 ADHD) because the above whole-brain image volumes were unavailable or because they had excessive head motion defined *a priori* as >2mm grand mean frame-to-frame displacement across all 6 tasks. After these restrictions, we included 175 total subjects (HC=76, SCZ=32, BPAD=35, ADHD=32).

### Neuropsychological testing and neurocognitive tasks

All individuals in the CNP dataset completed extensive neuropsychologic and neurocognitive testing, which have been thoroughly described in Poldrack (Poldrack et al., 2016). This testing took place outside of the MRI scanner. We aimed to predict an individual’s performance within five cognitive constructs: working memory, short-term memory, long-term memory, intelligence quotient (I.Q.), and executive function. Specific neuropsychologic and neurocognitive measures included in each phenotype may be referenced in Supplementary Figure 8. All measures used were available for all 175 included subjects.

#### CNP imaging parameters and preprocessing

Details of the image acquisition parameters have been published elsewhere. (Poldrack et al., 2016) In brief, all data were acquired on one of two 3T Siemens Trio scanners at UCLA. Functional MRI data were collected using a T2*-weighted echoplanar imaging (EPI) sequence with the following parameters: slice thickness=4mm, 34 slices, TR=2s, TE=30ms, flip angle=90°, matrix 64×64, FOV=192mm, oblique slice orientation. High-resolution anatomical MPRAGE data were also collected with the following parameters: TR=1.9s, TE=2.26ms, FOV=250mm, matrix=256×256, sagittal plane, slice thickness = 1 mm, 176 slices. From this dataset, we used the six task-based fMRI experiments described above.

All subsequent preprocessing was performed using image analysis tools available in BioImage Suite (Joshi et al., n.d.) and included standard preprocessing procedures (Finn et al., 2015), including removal of motion-related components of the signal; regression of mean time courses in white matter, cerebrospinal fluid, and gray matter; removal of the linear trend; and low-pass filtering(Greene et al., 2018). Mean frame-to-frame displacement yielded nine motion values per subject; these were used for subject exclusion and motion analyses. All subsequent analyses and visualizations were performed in BioImage Suite (Joshi et al., n.d.), Matlab (Mathworks), and Python.

#### Functional parcellation and network definition

We used the Shen 268-node atlas to parcellate the fMRI data into functionally coherent nodes (Shen, Tokoglu, Papademetris, & Constable, 2013). The Shen 268-node atlas is derived from an independent data set using a group-wise spectral clustering algorithm. The mean time courses of each node pair were correlated and correlation coefficients were Fisher transformed, generating six 268 × 268 connectivity matrices per subject for each fMRI task (see Figure 1.B); therefore, accounting redundant connections, each individual had a total of 214,668 edges.

**Figure 1.**
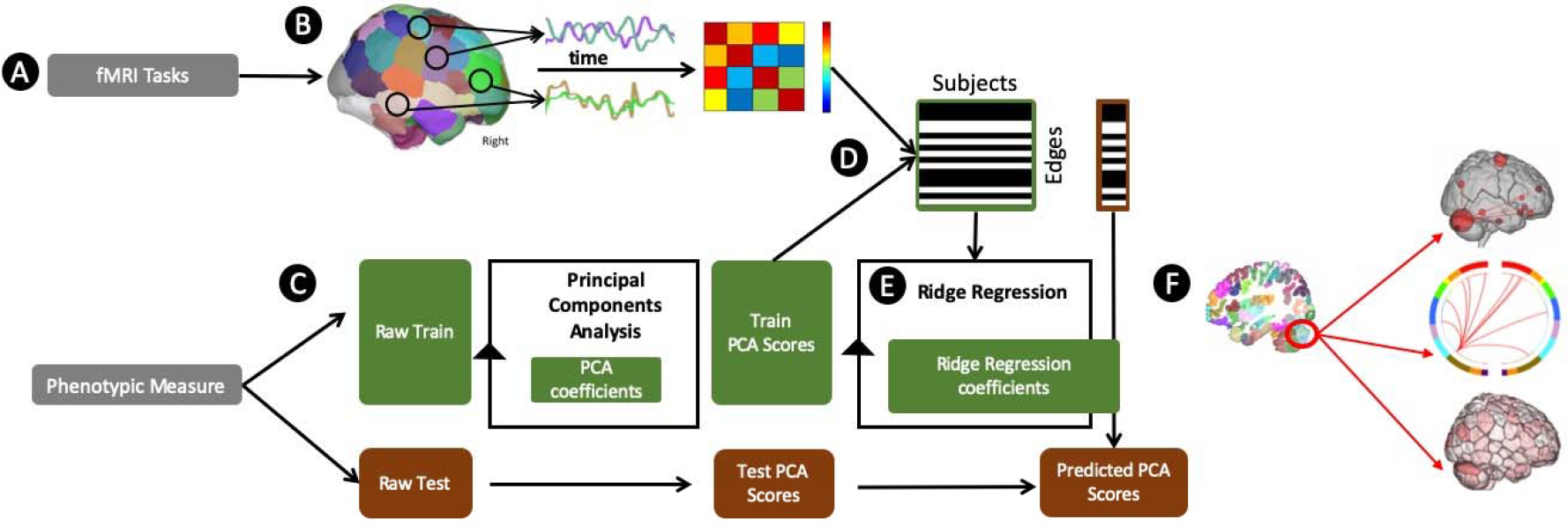
Overview of processing pipeline. (A) We used 6 fMRI tasks and 5 categories of phenotypic measures from the Neuropsychiatric Phenomics Consortium dataset (see Methods and Supplementary Materials). (B) We preprocessed and divided fMRI volumes using the Shen 268 node atlas. We then created a cross-correlation matrix of inter-node connectivity, hereafter described as edges. (C) We separated the behavioral and (D) fMRI data into train and test groups. We performed a principle components analysis to summarize one behavioral construct score per subject; we used the training data’s PCA coefficients to transform the behavioral test data into component space. (D) Across training subjects, we correlated each edge to phenotypic scores and restricted subsequent analyses to only significantly correlated edges (see Supplementary Materials for different statistical thresholds). (E) We used a ridge regression algorithm to train a predictive model wherein edges from all 6 fMRI tasks predict a phenotypic score. We applied this model to predict phenotypic score for each individual in the test group. (F) We used multiple types of illustrations to visualize the model structure. Here, we illustrate weighted edges from a cerebellar node as (from top-to-bottom): a node-and-edge plot wherein weighted edges are shown with their respective nodes and node size represents degree (or how many edges were weighted per node); a circle plot wherein cerebellar edges are connected to corresponding networks; and surface degree plots wherein node color indicates the node degree (darker colors are higher degree). Model performance measures are described in Methods.

The same spectral clustering algorithm was used to assign these 268 nodes to 8 networks (Finn et al., 2015; Shen et al., 2017), and the subcortical-cerebellar network was split into networks 8–10 (Noble et al., 2017). These networks are named based on their approximate correspondence to previously defined resting-state networks, and are numbered for convenience according to the following scheme: 1. Medial frontal, 2. Frontoparietal, 3. Default mode, 4. Motor, 5. Visual A, 6. Visual B, 7. Visual association, 8. Cingulo-Opercular, 9. Subcortical, 10. Cerebellum.

#### Principal Components Analysis

Given that a single behavioral measurement incompletely approximates a behavioral construct and has substantial noise due to individual variability and test (administration) variability, we summarized across multiple individual measures within each construct using principal components analysis to create what we consider a phenotypic measure. A similar strategy has been successfully employed to create a phenotypic measure of intelligence across individual measures of crystallized ability, processing speed, visuospatial ability and memory (Dubois et al., 2018). To maintain separate train and test groups, for each iteration, each PCA was limited to training datasets and these PCA coefficients were subsequently applied to the test dataset (see Figure 1.C).

#### CPM Ridge Regression

In ordinary least-squares (OLS) regression, a greater number of independent variables compared to the number of observations leads to an ill-posed problem and overfitting. To solve this ill-posed problem, regularization on regression coefficients can be applied to shrink the coefficients. Ridge regression shrinks the regression coefficients by imposing a *L2*-norm penalty on their size. Compared with OLS regression, the coefficients from ridge regression minimize a penalized residual sum of squares,

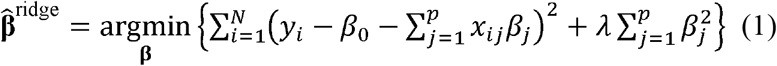

where *λ* is the complexity parameter that controls the shrinkage strength: *λ* = 0 *gives rise to the unregularized OLS, while* increasing *λ* shrinks the coefficients towards zero. If we write the criterion in equation (1) in matrix form, RSS(λ) = (**y** − **Xβ**)^T^(**y** − **Xβ**) + λ**β**^T^**β**, the ridge regression solutions can be solved by

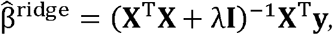

*where **I** is the p × p identity matrix. Compared with the solution for OLS*, 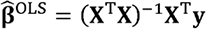, *adding a positive constant to the diagonal of* **X**^T^**X** *before inversion makes the problem nonsingular, even if* **X**^T^**X** *is not of full rank.*

Based on ridge regression, we modified the original CPM framework(Shen et al., 2017) to better suit the high-dimensional nature of connectivity data. In particular, we were ab (Figure 1). Specifically, due to the positive semi-definite nature of a functional connectivity matrix, the edges are not independent. Ridge regression is more robust than OLS in this case.

Instead of summing selected edges and fitting a one-dimensional OLS model, we directly fit a ridge regression model using the selected edges from all the tasks and apply the model to novel subjects in the cross-validation framework. *λ* parameter in the ridge regression is chosen by 10-fold cross-validation using only the training subjects. The largest *λ* value that has a mean squared error (MSE) within one standard error of the minimum MSE is chosen.

### Mass Multivariate Analysis, Theoretical Overview

In this section we recap a number of variance analysis techniques. Let’s assume *μ*_l_ and *μ*_2_ are means of two groups *g*_1_ and *g*_2_. A t-test aims to find if both means coming from same distribution of means like 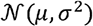. A univariate analysis of variance (ANOVA) aims to address whether groups *g*_1_, *g*_2_, ‥, *g*_m_ with means *μ*_l_, *μ*_2_,‥, *μ*_m_ where m > 2 come from same distribution 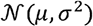. In this paper we focus on a multivariate equivalent of a t-test which is multivariate analysis of variance (MANOVA). A MANOVA is a multi-variate equivalent of t-test where we have vectors of means and the goal is to find out if they are sampled from a same distribution: 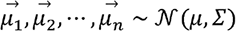 Therefore, a MANOVA gives the probability of sampling these vectors from 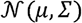. In other words, a MANOVA runs an overall test if vectors of means for different groups for a single variable are same or not. In a binary example, for variable *x*_*j*_, MANOVA tests if groups *x*_*j*_ =1 and *x*_*j*_ =0 have same mean vectors [*μ*_l_, *μ*_2_, *μ*_3_] corresponding to outcome variables (*y*_1_, *y*_2_, *y*_3_).

### Hypothesis Testing in MANOVA (Carey:1998) [supplementary]

We define the within group observed covariance matrix by 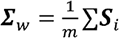.

Let’s assume 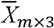 is a matrix that 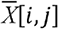 indicates the mean for *i*-th group on the *j*-th variable then estimation for *Σ* based on these *m* means is:

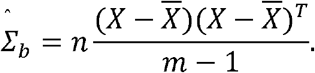

Similar to ANOVA we expect 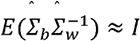.

Let’s assume each group has a normal distribution with covariance matrix *Σ* and the average between the groups is also the same estimation. An alternative hypothesis addresses the covariance matrix on sampling distribution of means coming from a normal distribution with mean *μ* and covariance matrix 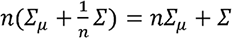.

Then we have the following equations:

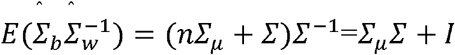

Exactly analogous to ANOVA, diagonal values of the results are greater than or equal to 1. There are different ways of analyzing this ratio including but not limited to Pillai’s trace 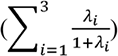, Hotelling-Lawley’s trace 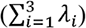, and Wilk’s lambda 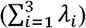 where 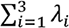 indicates *i*-th eigenvalue of the ratio.

To correct for multiple comparisons, we used a network based statistic (NBS) with 1000 iterations. Edges were considered significant if they appear in a cluster larger than a null distribution of clusters, as defined in the permutation testing. In other words significant edges are more likely to provoke nearest neighbors to be significant.

### Mass Multivariate Analysis, Disease Group Differences

To assess for statistically significant differences in connectomes based on symptom-based disease categories, we performed a mass multi-variate analysis on separate disease groups, as defined by the DSM-IV (First et al., 2002) by the UCLA group (Poldrack et al., 2016). These groups were healthy controls, schizophrenia, bipolar disorder, and ADHD.

Figure 4 illustrates group differences among all four disease groups with surface plots. We use this as a mask for the next experiments. First we break down it into all six possible bi-groups and plot edge-wise connections only those span on the initial mask. Our results show only 368 edges remaining significant after NBS correction. Last row show heat maps in network level. Indeed, we aggregate edges based on their membership to 10 different networks and see if there is meaningful connections between them. We used normalization based on network size since each network consisting of different number of nodes.

### Mass Multivariate Analysis, Disease Severity within Medication Class

To assess for statistically significant differences in connectomes based on disease severity, we performed a mass multivariate analysis on each medication class as a function of symptomatic remission. (Note that a two group MANOVA will give the same results as Hotelling’s T2.)

We grouped patients based on whether they were prescribed an antipsychotic (n=46), mood stabilizer (n=24), or antidepressant (n=31; See Supplementary Figure 10). Given the limited sample size, we combined mood stabilizers and antidepressants to create two medication class: psychotic and affective disorder medications.

We defined remission based on standard symptom-based criteria. For affective medications, we defined remission as a Hamilton 28-item Depression scale (HAMD-28) of ≤10 (Hamilton, 1960). For psychosis medications, we defined remission (Alonso, Ciudad, Casado, & Gilaberte, 2008) as a Scale for Assessment of Positive Symptoms (SAPS), ratings of mild or less (≤2) for delusions, hallucinations, positive formal thought disorder, and bizarre behavior considered individually. For the Scale for Assessment of Negative Symptoms (SANS), ratings of mild or less (≤2) for affective flattening, avolition-apathy, anhedonia-asociality, alogia considered individually.

### Statistical Analyses

#### Cross-validation and performance measures

We used two types of cross-validation methods: 10-fold cross validation (see Figure 2) and leave-one-group out (i.e. one clinical group; see Figure 3).

**Figure 2.**
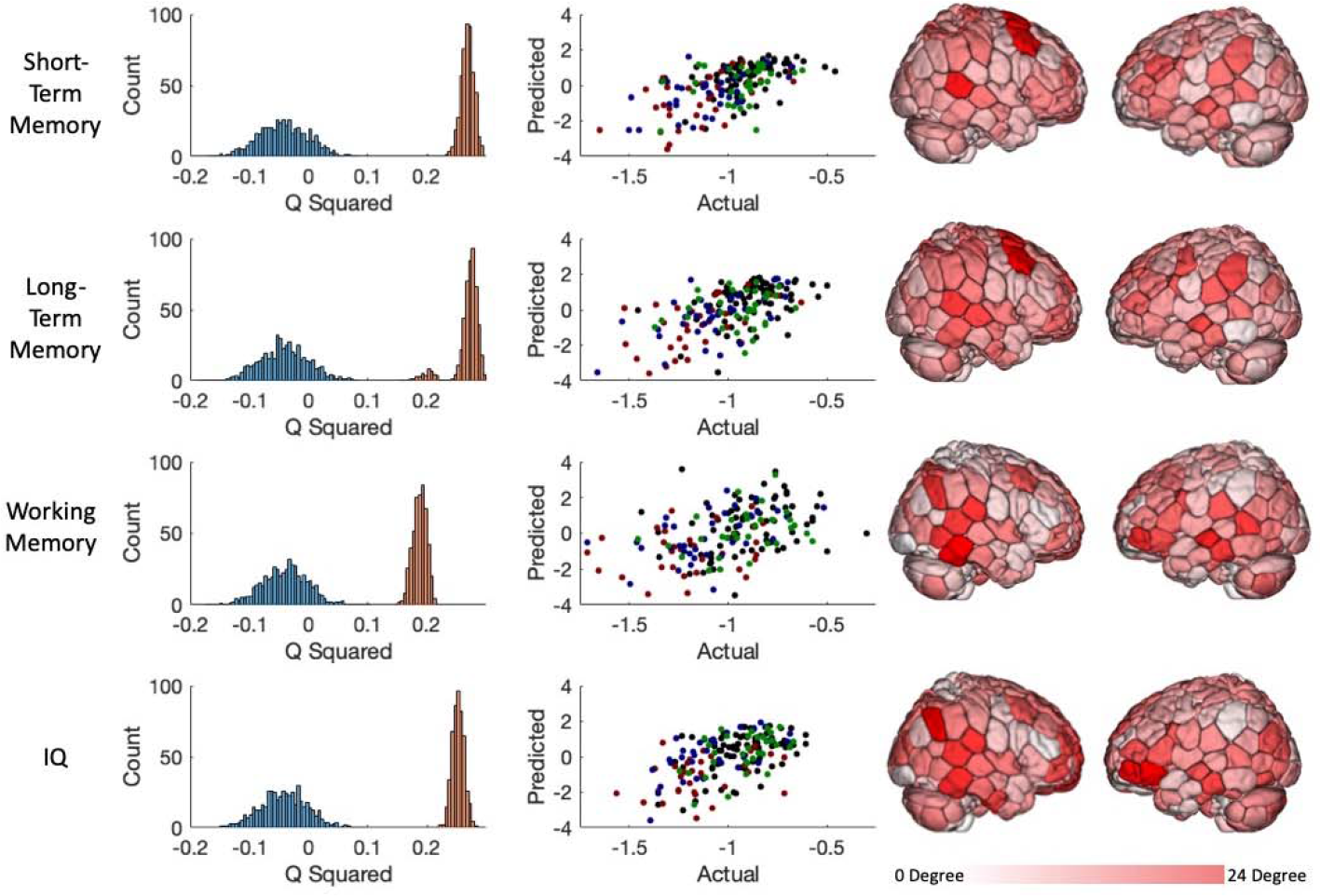
10-fold phenotype predictive model performance. The left column shows a histogram of the model performance across 1000 iterations of the actual (red) and randomly permuted (blue) data. The middle column shows how actual and predicted values compare for the median-performing model (actual distribution in left column; black dots are controls; green, schizophrenia; blue, bipolar disorder; red, ADHD). The right columns shows surface plots of each node’s degree (the number of weighted edges per node).

**Figure 3.**
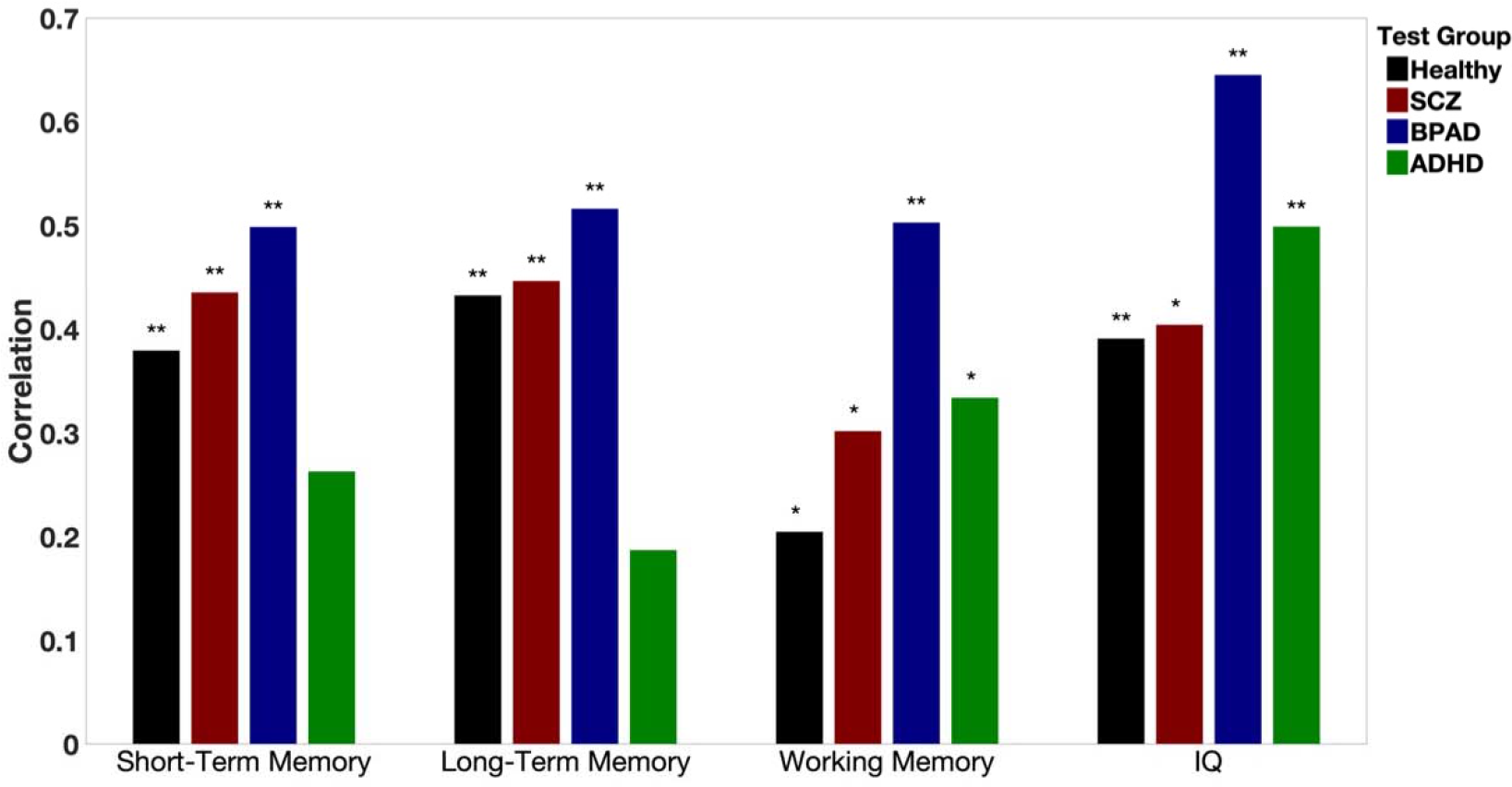
Leave one group out phenotype predictive model performance. Each transdiagnostic model was trained on three of the clinical groups and tested on the clinical group indicated below (i.e. performance for the models tested on. “Healthy” were trained on “SCZ”, “BRAD”, and “ADHD” data). Performance measured as the Pearson correlation between actual and predicted cognitive measure; ** indicates correlation significant at p<0.01 for one-tail; * indicates p<0.05. SCZ=schizophrenia; BPAD=bipolar affective disorder.

In the 10-fold cross-validation, the sample was randomly divided into 10, approximately equal-sized groups; on each fold, the model was trained on 9 groups and tested on the excluded 10th group. This process was repeated iteratively, with each group excluded once. Unless otherwise specified (cf. the Supplementary Materials), for each 10-fold analysis, we create random divisions from the dataset over 1,000 iterations. CPM was performed with a range of edge selection thresholds from 0.001 to 0.05, which did not substantially change model performance (see Supplementary Figure 8). Model performance was evaluated by the cross-validated *R*^*2*^,

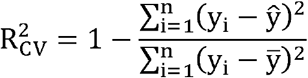

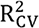 can be negative (Scheinost et al., 2019) and negative values were set to 0. In this paper, 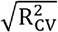 is reported as it is comparable to, but less biased than, the normally used Pearson correlation value when using cross-validation. 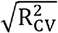 is averaged over the cross validation folds.

Because the leave-one-group out analyses exhausted all possible combinations of groups, we did not iterate these analyses. Because a separate model was trained and evaluated for each subject group, we used a Pearson correlation between actual and predicted cognitive phenotype for each test group to evaluate model performance.

#### Node Contribution to Ridge Regression Model

Because each ridge regression model represents a linear combination across (in some cases) thousands of edges (see Supplementary Figures), we are unable to fully illustrate edge contribution in both a mathematically complete and cognitively interpretable way. To best summarize which edges the ridge regression algorithm weighted most, we created two types of illustrations: 1) a brain surface plot showing each edge that was significantly correlated with the behavioral constructs in the training group (Weighted node contribution = Edge1weight\**sd*(Edge1weight across all subjects), where *sd* stands for the standard deviation across iterations; 2) a circle plot wherein network edges are plotted between the networks with which they interact.

#### Statistical thresholds

For the mass multivariate connectomic analyses, we corrected all statistical analyses for multiple comparisons using the Network-Based Statistic False Discovery Rate correction (NBS-FDR) at p<0.05.

#### Effects of motion

Motion is an important confound for estimates of functional connectivity. In our analysis, motion was significantly correlated with most individual behavioral measures (see supplementary materials). We used two accepted methods for accounting for this confound: we excluded all subjects with >0.2 mean frame-to-frame motion across all tasks (Hsu, Rosenberg, Scheinost, Constable, & Chun, 2018) and we regressed out motion during the feature-selection step (see Figure 1) using partial correlation (i.e. as opposed to correlation). All results presented in the main text were performed using the partial correlation step, however this did not substantially change model performance (see Supplementary Figure 6).

#### Code availability

Matlab scripts to run the main CPM analyses can be found at (https://www.nitrc.org/projects/bioimagesuite/). BioImage Suite tools used for analysis and visualization can be accessed at (http://bisweb.yale.edu). Matlab scripts written to perform additional post-hoc analyses are available from the authors upon request. The median model (i.e. in terms of performance across 1,000 iterations of the 10-fold analysis) for our connectome-based predictive model is also available at the BioImage Suite. Access to the web-state is available at: http://bisweb.yale.edu/XXXXX

#### Network overlap

To explicitly explore the macroscale brain networks that were predictive of each behavioral construct (i.e. in the 10-fold predictive model) or were significantly different across disease groups or treatment response in affective or psychotic medications (i.e. in the mass-multivariate analyses), we assigned each selected (in predictive models) or significantly different (in mass-multivariate analyses) edge to a pair of canonical networks such that edge (*i,j*) would be assigned to the network that includes node *i* and the network that includes node *j*. For the 10-fold predictive analysis, an edge was considered “weighted” if it was weighted in all of the 10 folds and in >95% of the iterations (>950 of 1000 iterations). For the mass multivariate analysis, an edge was considered “significant” if it passed the statistical threshold described above.

Network overlap was determined with the hypergeometric cumulative density function, which returns the probability of drawing up to *x* of *K* possible items in *n* drawings without replacement from an *M*-item population. This was implemented in Matlab as follows: *p=1-hygecdf(x, M, K, n)* where *x* equals the number of weighted/significant edges, *n* equals the total number of weighted/significant edges in the brain, *K* equals the total number of edges (whether weighted/significant or not) in the network of interest, and *M* equals the total number of edges (whether weighted/significant or not) in the brain.

## RESULTS

### Phenotype 10-Fold predictive model performance

Across all behavioral constructs, we were able to significantly predict performance in a transdiagnostic fashion. The 6 task-based fMRI connectomes successfully predicted short-term memory, long-term memory, and working memory and IQ, but did not predict executive function. Successful prediction results are shown in Figure 2; executive function are shown in Supplementary Materials. It is possible that executive function prediction failed because we were unable to summarize the variance across the multiple executive function measures in a single component (see Supplementary Figure 8.E). While in general task contribution to predictive performance across tasks was uniform, we noticed that for short and long-term memory, the PAM-RET and BART tasks contributed the most to overall prediction; for IQ prediction, the PAM-RET task contributed the most (see Supplementary Figure 7). Across diagnoses, the population distributions overlapped for both actual and predicted phenotypes (See Figure 2, middle column).

In line with previous CPM results, our models were complex and span the entire brain. We conducted multiple complementary follow-up analyses to assess the robustness of our results. We tested the effect of sample size, motion, number of edges, and edge selection statistical threshold on model performance. These analyses are found in Supplementary Materials.

### Phenotype leave one group out predictive model performance

We were able to significantly predict performance across groups. In 14 of 16 analyses, models trained in all but one group were able to successfully predict the measure of interest in the left-out group. This was true even when models were trained only on patients and tested on healthy controls. These results were robust to the effect of edge selection threshold, which may be found in the Supplementary Figures.

### Mass Multivariate Analysis, Diagnostic Category Differences

We found significant differences between each clinical group’s task-based functional connectomes, indicating that while connectomes were able to predict cognitive performance across diagnosis, these connectomes had significant differences. Only 368 edges passed our strict network-based correction for multiple comparison, representing only 0.17% of possible edges. Surface plots showing group differences among all four disease groups are shown in Figure 4. The spatial locations of the edges is described in Figure 6.

**Figure 4.**
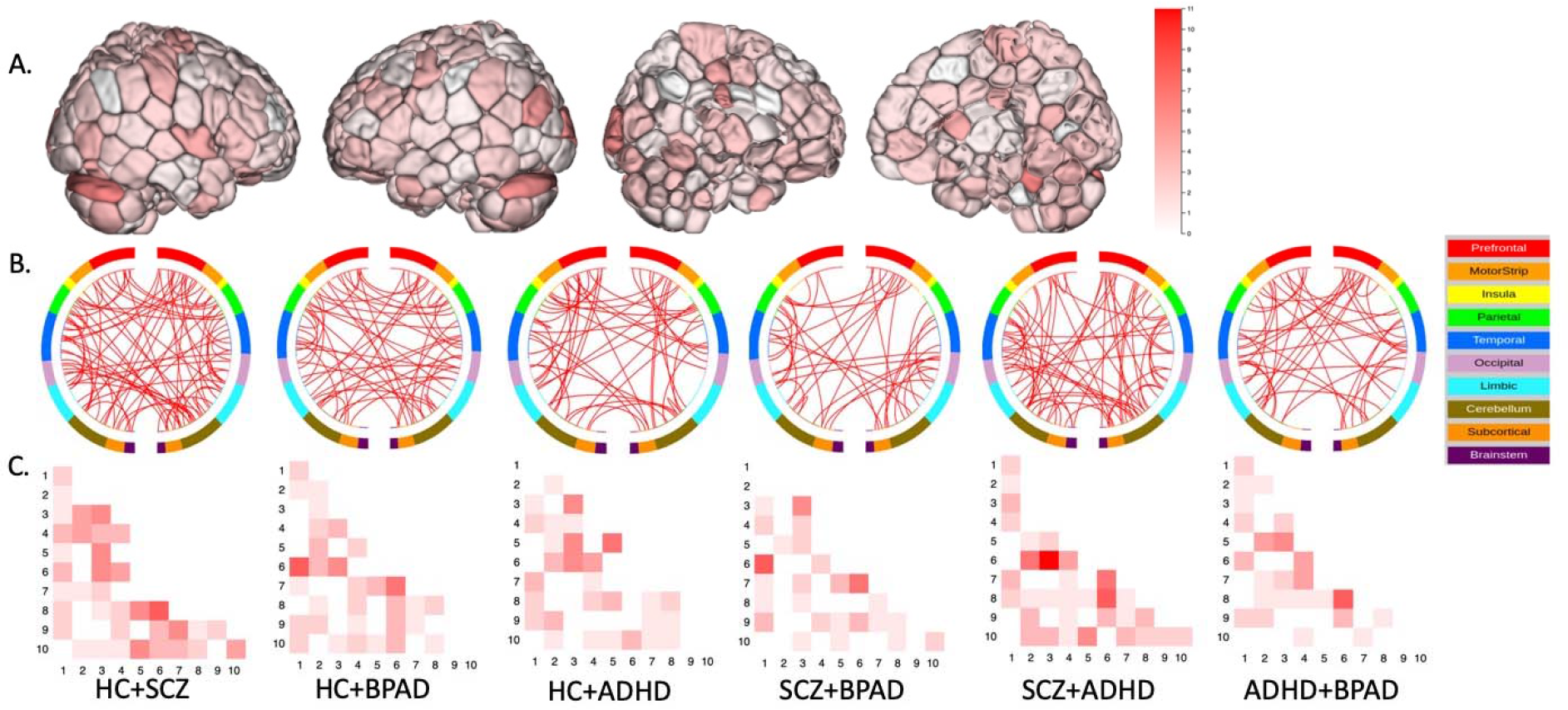
Mass multivariate analysis of disease group difference in brain network structure across all tasks. (A) Surface illustration of nodes where edges (network connections) significantly differ across all clinical groups, as measured with Hotelling’s T2. (B) and (C) illustrate significantly different edges across the two indicated groups. (B) illustrates circle plots that were not thresholded by degree while (C) illustrates network-to-network edge interactions with the most significant connections. (Network Labels: 1 = medial frontal, 2 = frontoparietal, 3 = default mode, 4 = motor cortex, 5 = visual A, 6 = visual B, 7 = visual association, 8 = salience, 9 = subcortical, 10 = cerebellum)

**Figure 5.**
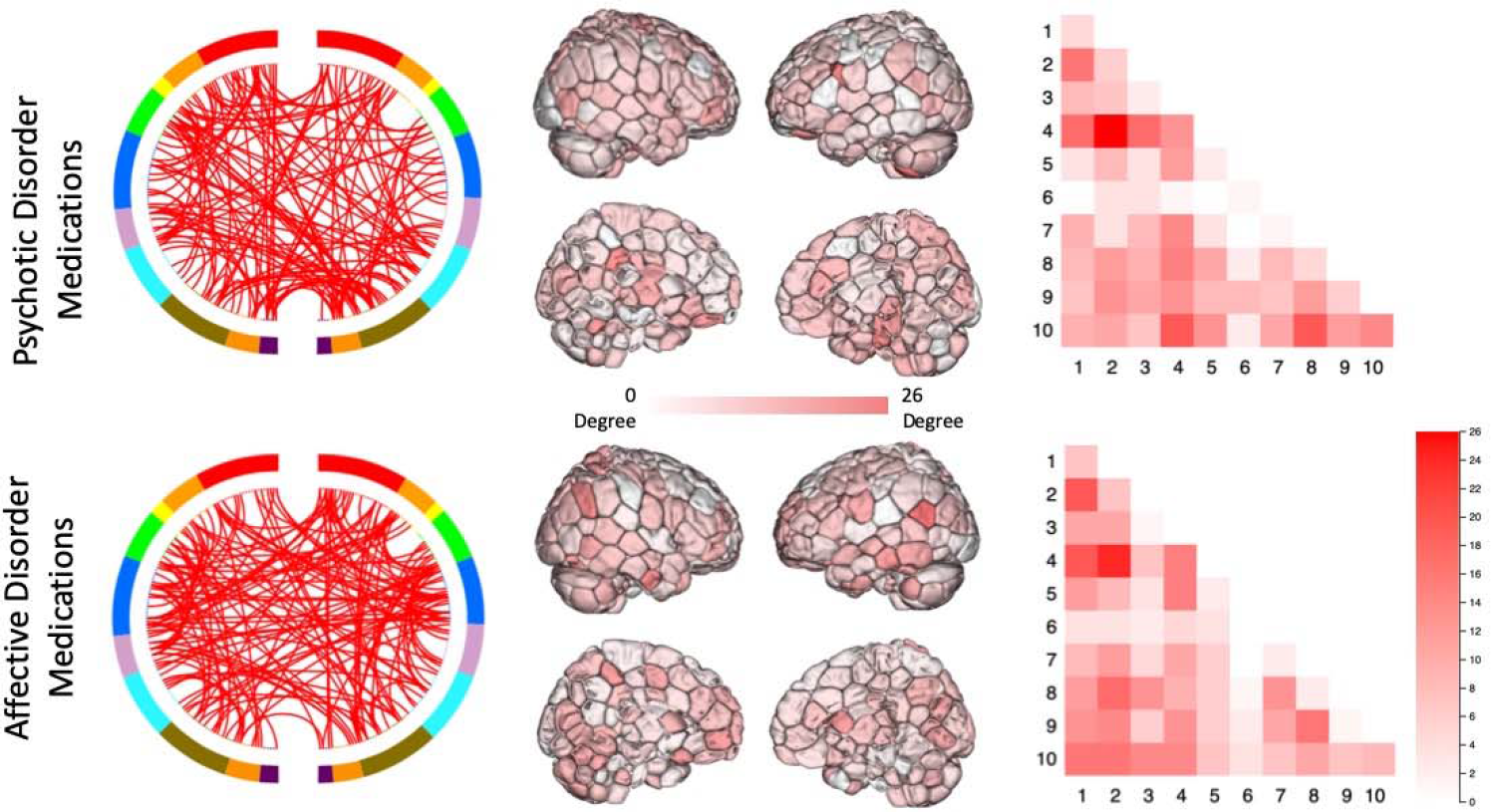
Mass multivariate analysis of disease severity by pharmacologic class. The top row evaluates all patients (across all diagnoses) who were prescribed an antipsychotic and illustrates where responders and non-responders (see Methods for definition) differed in terms of: (left) circle plots of significantly different edges, (middle) surface plots of node degree where edges (network connections) significantly differed, (right) network-based plots showing how many edges significantly differed by network hubs. (Network Labels: 1= medial frontal, 2 = frontoparietal, 3 = default mode, 4 = motor cortex, 5 = visual A, 6 = visual B, 7 = visual association, 8 = salience, 9 = subcortical, 10 = cerebellum)

**Figure 6.**
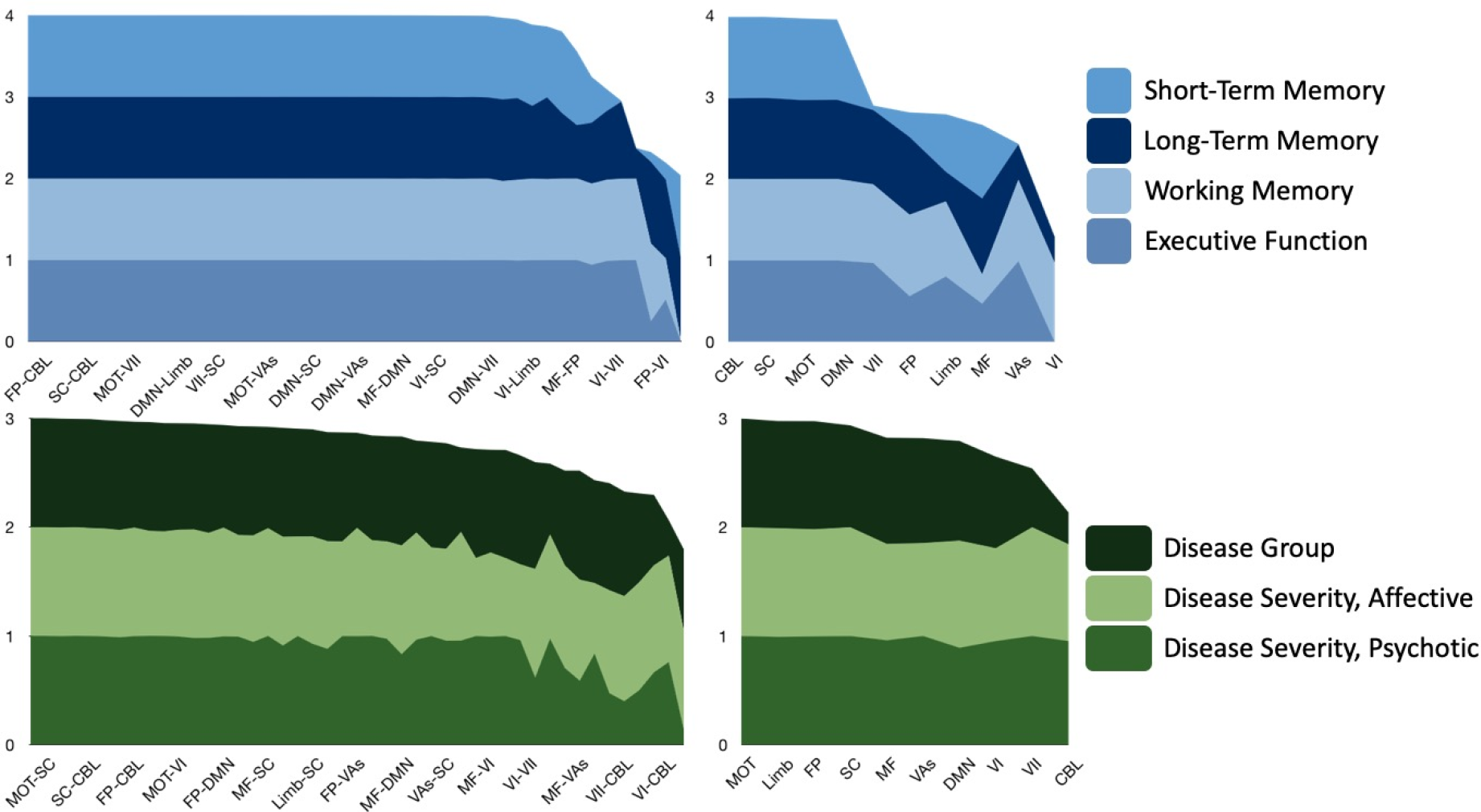
Network weightings in predictive (above) and mass multivariate (below) analyses. Layer thickness represents the likelihood that a particular inter-node (left column) or intra-node (right column) edge was selected by the model, as computed by the hypergeometric distribution.

### Mass Multivariate Analysis, Disease Severity within Medication Class

We found significant differences between remitters and non-remitters in patients taking medications for both psychotic and affective disorders, as shown in Figure 5. These differences persisted even after a conservative correction for multiple corrections (p<0.05 FWER-corrected via NBS). The psychotic disorder medication class showed only 405 significant edges (0.019% of possible edges), while the affective disorder medication class showed 415 (0.019% of possible edges). Significantly different edges were distributed throughout the brain, however it appears that the motor and cerebellar networks had the highest density of edges, as further explored below and in Figure 6.

### Network Profile and Overlap

We found similar network trends across predictive models of behavioral constructs, as shown in Figure 6. These trends were also similar in the mass multivariate analyses of disease class and disease severity. Edges within the cerebellar, fronto-parietal, default mode, subcortical, and motor networks were more likely (i.e. according to the hypergeometric distribution, see Methods) to be highly weighted across the 10-fold predictive analyses. In turn edges within these same areas were more likely to be significantly different than others in the brain.

## DISCUSSION

The unifying goal across our analyses is to evaluate how different frameworks capture the brain basis of deficits seen in psychiatric disease. Our work suggests that the same macroscale brain circuitry underlies a given cognitive function across all people, regardless of the diagnoses they carry, and that apparently distinct behavioral endpoints may be caused by individual differences within these common circuits. We further show that three common frameworks for characterizing clinical deficits (phenotype, disease group, and disease severity) each offer useful and perhaps complimentary explanations for the brain’s underlying functional architecture as affected by psychiatric disease.

### Brain basis of disease

Although patients with mental illness have historically been binned into symptom-based categories, we present evidence that this is not the only neurobiologically valid way of viewing phenotypic variability within and across patients. At least on the four phenotypes we evaluated (short-term memory, long-term memory, working memory, and executive function), we show that models built from patients diagnosed with schizophrenia, bipolar disorder, and ADHD can predict behavioral phenotypes in so-called healthy controls. This indicates that neural representation underlying a given behavioral phenotype remains the same notwithstanding disease category or symptom burden.

We also show (in the same dataset) statistically significant differences in the brain’s functional organization based on disease group and disease severity. Our disease group mass multivariate analysis indicated statistically significant differences in similar brain networks identified by our phenotype prediction models (see Figure 6). Furthermore, when we collapse across disease group to form two medication classes, we showed that measures of disease severity also identify statistically significant differences in the brain’s functional organization. Therefore, our results suggest that each framework—phenotype, disease group, and disease severity—has a neurobiologic correlate.

The ultimate question, then, is which framework is most clinically useful. How can we most ably define clinical deficits in such a way that they can be identified and alleviated? Data-driven methods that account for, characterize, and cluster (where necessary) disease-relevant phenotypic variance increasingly appear to be the way forward; especially in studies that seek to detect some neurobiologic signal for mental illness. Perhaps here, a comparison can be made to clinical decision: the question isn’t which *disease group* a patient falls into, but rather how to best predict which treatment will alleviate a patient’s symptom pattern (Chekroud et al., 2017; Drysdale et al., 2017). Although this is intuitive to clinicians—who hope to modify a patient’s phenotype or disease severity, not their disease group—this is not how researchers have traditionally analyzed patient data.

### Complex Model Reduction

We have struggled to reduce the results across 214,668 edges to something that is cognitively manageable—at minimum by us, the authors. This is an important insight into biomarker development: useful models of brain-behavior are not necessarily simple (Rosenberg, Casey, & Holmes, 2018). Just as an overly simplistic model cannot be compensated for by an increased sample size (underfitting), an overly complex model will not necessarily generalize (overfitting). Although communicating a complex model (i.e. in a paper or figure or conversation) requires a reduction of complexity, this can appear to come at the cost of learning something neurobiolgically meaningful. There isn’t necessarily a simple answer to any neurobiologic question. What we’re learning about the brain and its functional connections is that it’s complex. The traditional function-location framework appears to be insufficient to understand clinically-relevant brain-behavior relationships.

Accordingly, we uploaded the complete predictive model (based on the median-performing iteration, see Figure 2) and have created a freely-accessible instantiation of Bioimage Suite online wherein readers may access and navigate the entire model [We’ll negotiate this with the journal].

### Cerebellum

Consistent with many previous reports, our predictive and explanatory models implicate the cerebellum as a network that sub-serves more than motor function. Functional neuroimaging evidence supports the parcellation of the cerebellum into at least three regions associated with sensorimotor, cognitive, and limbic functions (Schmahmann & Caplan, 2006) (Riedel et al., 2015). We report that the cerebellum is influential in predicting phenotypic traits and in explaining symptomatic remission within pharmacologic classes. Likewise, our data further suggests that even cerebral “motor” areas (including motor, somatosensory, primary auditory cortices) also support more than motor function. Our results add to the growing literature that brain regions traditionally defined as “motor” influence phenotypic traits, symptom profile, and treatment response (cf. Figures 2, 3, 4).

## REFERENCES

Alonso, J., Ciudad, A., Casado, A., & Gilaberte, I. (2008). Measuring schizophrenia remission in clinical practice. Canadian Journal of Psychiatry. Revue Canadienne De Psychiatrie, 53(3), 202–206. http://doi.org/10.1177/070674370805300311

Barron, D. S. (2019). Should Mental Disorders Have Names? Retrieved May 15, 2019, from

Barron, D. S., Salehi, M., Browning, M., Harmer, C. J., Constable, R. T., & Duff, E. (2018). Exploring the prediction of emotional valence and pharmacologic effect across fMRI studies of antidepressants. NeuroImage: Clinical, 20, 407–414. http://doi.org/10.1016/j.nicl.2018.08.016

Chekroud, A. M., Gueorguieva, R., Krumholz, H. M., Trivedi, M. H., Krystal, J. H., & McCarthy, G. (2017). Reevaluating the Efficacy and Predictability of Antidepressant Treatments. JAMA Psychiatry, 74(4), 370–9. http://doi.org/10.1001/jamapsychiatry.2017.0025

Cross-Disorder Group of the Psychiatric Genomics Consortium. (2013). Identification of risk loci with shared effects on five major psychiatric disorders: a genome-wide analysis. Lancet (London, England), 381(9875), 1371–1379. http://doi.org/10.1016/S0140-6736(12)62129-1

Drysdale, A. T., Grosenick, L., Downar, J., Dunlop, K., Mansouri, F., Meng, Y., et al. (2017). Resting-state connectivity biomarkers define neurophysiological subtypes of depression. Nature Publishing Group, 23(1), 28–38. http://doi.org/10.1038/nm.4246

Dubois, J., Galdi, P., Paul, L. K., & Adolphs, R. (2018). A distributed brain network predicts general intelligence from resting-state human neuroimaging data. Philosophical Transactions of the Royal Society of London. Series B, Biological Sciences, 373(1756). http://doi.org/10.1098/rstb.2017.0284

Finn, E. S., Shen, X., Scheinost, D., Rosenberg, M. D., Huang, J., Chun, M. M., et al. (2015). Functional connectome fingerprinting: identifying individuals using patterns of brain connectivity. Nature Publishing Group, 18(11), 1664–1671. http://doi.org/10.1038/nn.4135

First, M. B., Spitzer, R. L., Gibbon, M., & Williams, J. (2002). Structured clinical interview for DSM-IV-TR axis I disorders, research version, patient edition.

Goodkind, M., Eickhoff, S. B., Oathes, D. J., Jiang, Y., Chang, A., Jones-Hagata, L. B., et al. (2015). Identification of a common neurobiological substrate for mental illness. JAMA Psychiatry, 72(4), 305–315. http://doi.org/10.1001/jamapsychiatry.2014.2206

Gorgolewski, K. J., Auer, T., Calhoun, V. D., Craddock, R. C., Das, S., Duff, E. P., et al. (2016). The brain imaging data structure, a format for organizing and describing outputs of neuroimaging experiments. Scientific Data, 3, 160044. http://doi.org/10.1038/sdata.2016.44

Greene, A. S., Gao, S., Scheinost, D., & Constable, R. T. (2018). Task-induced brain state manipulation improves prediction of individual traits. Nature Communications, 9(1), 2807. http://doi.org/10.1038/s41467-018-04920-3

Hamilton, M. (1960). A rating scale for depression. Journal of Neurology, Neurosurgery & Psychiatry, 23, 56–62. http://doi.org/10.1136/jnnp.23.1.56

Holmes, A. J., & Patrick, L. M. (2018). The Myth of Optimality in Clinical Neuroscience. Trends in Cognitive Sciences, 22(3), 241–257. http://doi.org/10.1016/j.tics.2017.12.006

Horien, C., Shen, X., Scheinost, D., & Constable, R. T. (2019). The individual functional connectome is unique and stable over months to years. NeuroImage, 189, 676–687. http://doi.org/10.1016/j.neuroimage.2019.02.002

Hsu, W.-T., Rosenberg, M. D., Scheinost, D., Constable, R. T., & Chun, M. M. (2018). Resting-state functional connectivity predicts neuroticism and extraversion in novel individuals. Social Cognitive and Affective Neuroscience, 13(2), 224–232. http://doi.org/10.1093/scan/nsy002

Insel, T. R., Sahakian, B. J., Voon, V., Nye, J., & Brown, V. (2012). Drug research: a plan for mental illness. Nature, 483(7389), 269–269. http://doi.org/10.1038/483269a

Insel, T., Cuthbert, B., Garvey, M., Heinssen, R., Pine, D. S., Quinn, K., et al. (2010). Research domain criteria (RDoC): toward a new classification framework for research on mental disorders. American Journal of Psychiatry, 167(7), 748–751. http://doi.org/10.1176/appi.ajp.2010.09091379

Joshi, A., Scheinost, D., Okuda, H., Belhachemi, D., Murphy, I., Staib, L. H., & Papademetris, X. (n.d.). Unified Framework for Development, Deployment and Robust Testing of Neuroimaging Algorithms. Neuroinformatics, 9(1), 69–84. http://doi.org/10.1007/s12021-010-9092-8

Noble, S., Spann, M. N., Tokoglu, F., Shen, X., Constable, R. T., & Scheinost, D. (2017). Influences on the Test-Retest Reliability of Functional Connectivity MRI and its Relationship with Behavioral Utility. Cerebral Cortex (New York, N.Y.: 1991), 27(11), 5415–5429. http://doi.org/10.1093/cercor/bhx230

Poldrack, R. A., Congdon, E., Triplett, W., Gorgolewski, K. J., Karlsgodt, K. H., Mumford, J. A., et al. (2016). A phenome-wide examination of neural and cognitive function. Scientific Data, 3, 160110. http://doi.org/10.1038/sdata.2016.110

Riedel, M. C., Ray, K. L., Dick, A. S., Sutherland, M. T., Hernandez, Z., Fox, P. M., et al. (2015). Meta-analytic connectivity and behavioral parcellation of the human cerebellum. NeuroImage, 117, 327–342. http://doi.org/10.1016/j.neuroimage.2015.05.008

Rosenberg, M. D., Casey, B. J., & Holmes, A. J. (2018). Prediction complements explanation in understanding the developing brain. Nature Communications, 1–13. http://doi.org/10.1038/s41467-018-02887-9

Rosenberg, M. D., Finn, E. S., Scheinost, D., Papademetris, X., Shen, X., Constable, R. T., & Chun, M. M. (2016). A neuromarker of sustained attention from whole-brain functional connectivity. Nature Neuroscience, 19(1), 165–171. http://doi.org/10.1038/nn.4179

Scheinost, D., Noble, S., Horien, C., Greene, A. S., Lake, E. M., Salehi, M., et al. (2019). Ten simple rules for predictive modeling of individual differences in neuroimaging. NeuroImage, 193, 35–45. http://doi.org/10.1016/j.neuroimage.2019.02.057

Schmahmann, J. D., & Caplan, D. (2006). Cognition, emotion and the cerebellum. Brain, 129(Pt 2), 290–292. http://doi.org/10.1093/brain/awh729

Shen, X., Finn, E. S., Scheinost, D., Rosenberg, M. D., Chun, M. M., Papademetris, X., & Constable, R. T. (2017). Using connectome-based predictive modeling to predict individual behavior from brain connectivity. Nature Publishing Group, 12(3), 506–518. http://doi.org/10.1038/nprot.2016.178

Shen, X., Tokoglu, F., Papademetris, X., & Constable, R. T. (2013). Groupwise whole-brain parcellation from resting-state fMRI data for network node identification. NeuroImage, 82(C), 403–415. http://doi.org/10.1016/j.neuroimage.2013.05.081

Vanasse, T. J., Fox, P. M., Barron, D. S., Robertson, M., Eickhoff, S. B., Lancaster, J. L., & Fox, P. T. (2018). BrainMap VBM: An environment for structural meta-analysis. Human Brain Mapping, 39(8), 3308–3325. http://doi.org/10.1002/hbm.24078

