## Supplementary Figures for "Task-Based Functional Connectomes Predict Cognitive Phenotypes Across Psychiatric Disease"

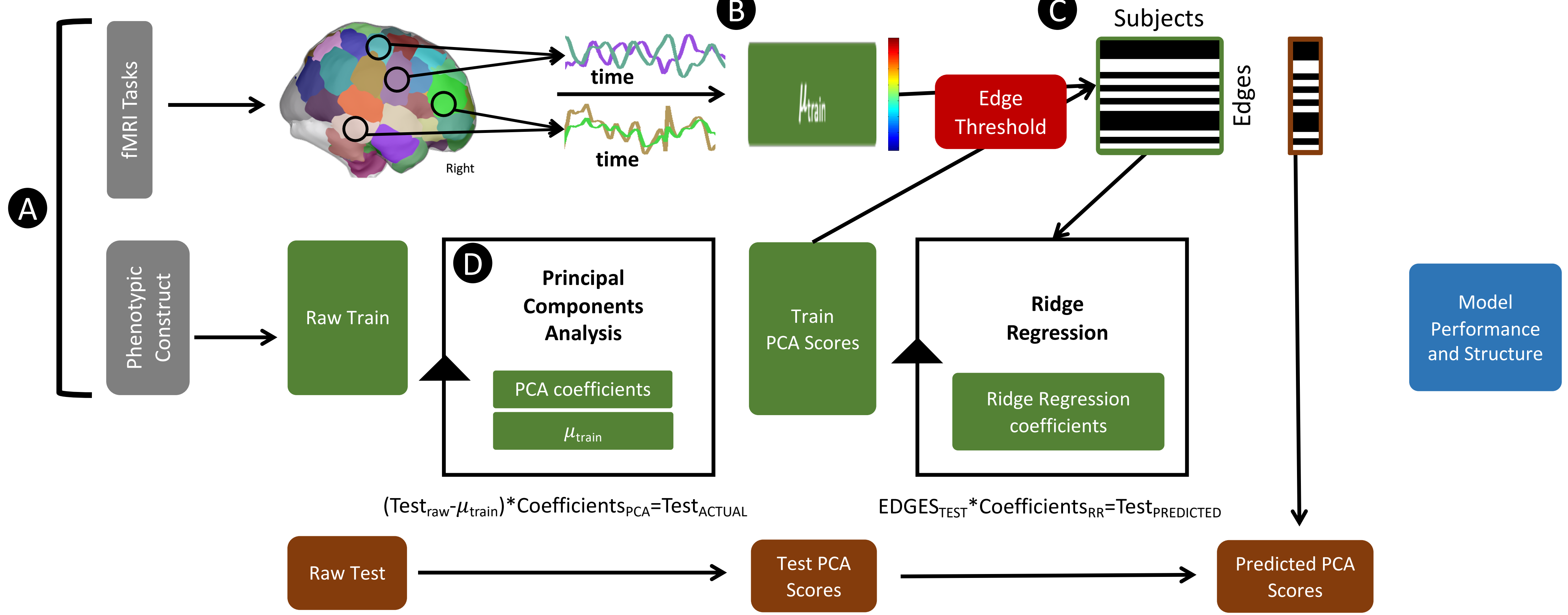


**Supplementary Figure 1. Overview of Supplementary Analyses**. (A) In Supplementary Figure 2, we show the effect of sample size on model performance for the 10-fold phenotype predictive analyses. (B) In Supplementary Figure 3, we show the effect of pre-threshold edge number (C.1) We tested the effect of different statistical significance thresholds on the correlation analysis, we ran the analyses at $p<0.01$, $p<0.005$, and $p<0.001$ thresholds for the k-10 (Figure 4) and leave-group-out (Figure 8) analyses. (C.2) We tested the effect of head motion on prediction performance by running analyses with and without motion regression (e.g. partial correlation, as reported in Figure 2). Results did not substantially change as shown in Supplementary Figure 6. D) We tested whether the different cognitive phenotypes could be adequately represented by a single (i.e. the first) principal component; this analysis showed that all constructs except for executive function can be (Supplementary Figure 9).

**Supplementary Figure 2.** Performance as a function of sample size for 10-fold ridge regression analysis. We iterated these analyses 200 times to produce the error bars. We did not regress out motion in these analyses.

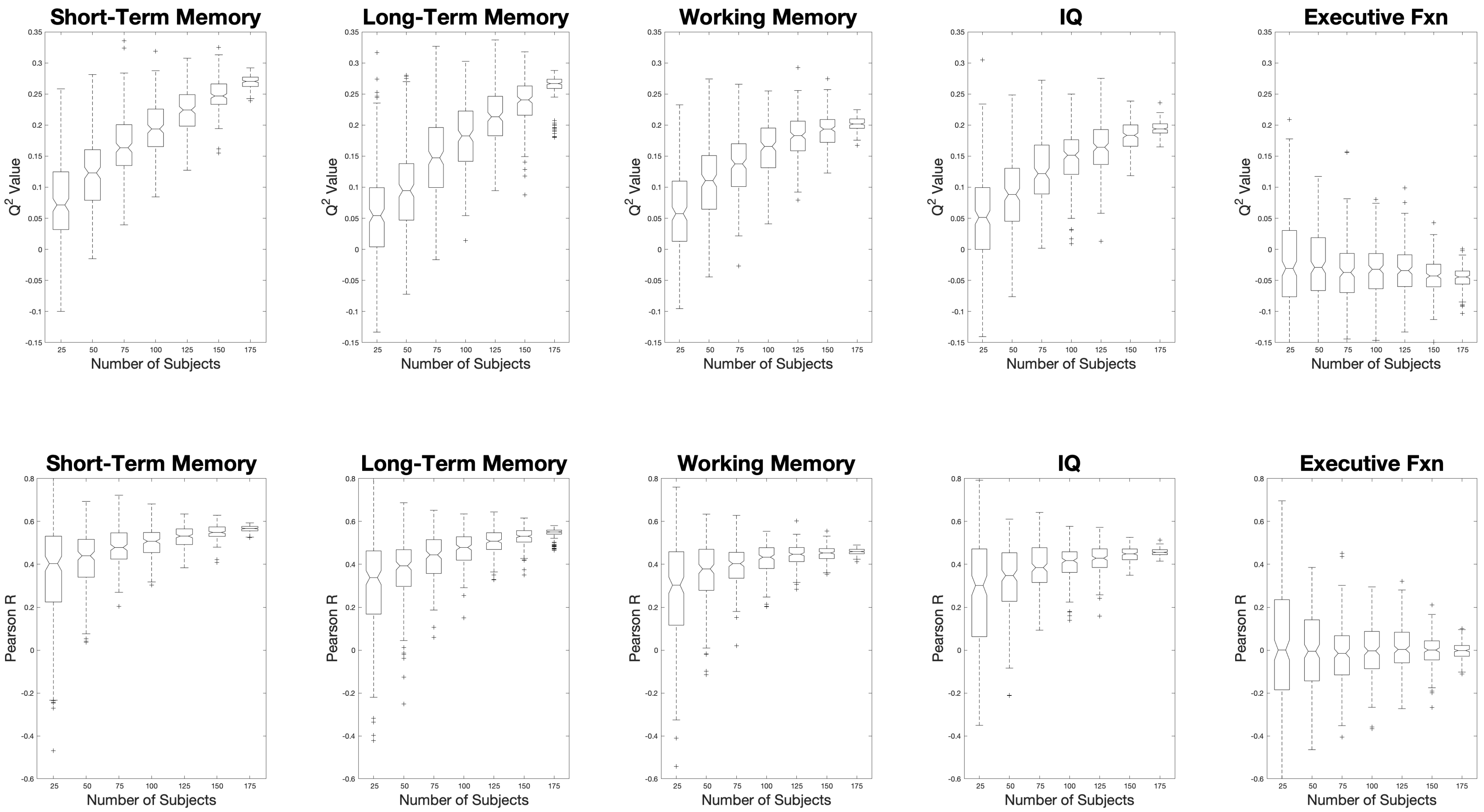

**Supplementary Figure 3.** Performance as a function of pre-thresholded edge number for 10-fold ridge regression analysis. All edge increments were randomly selected from the total number. Top row shows performance measured with Q-Squared. Bottom Row shows performance measured as Pearson correlation between actual and predicted phenotype scores. We iterated analyses 200 times to produce error bars; we did not regress out motion in these analyses.

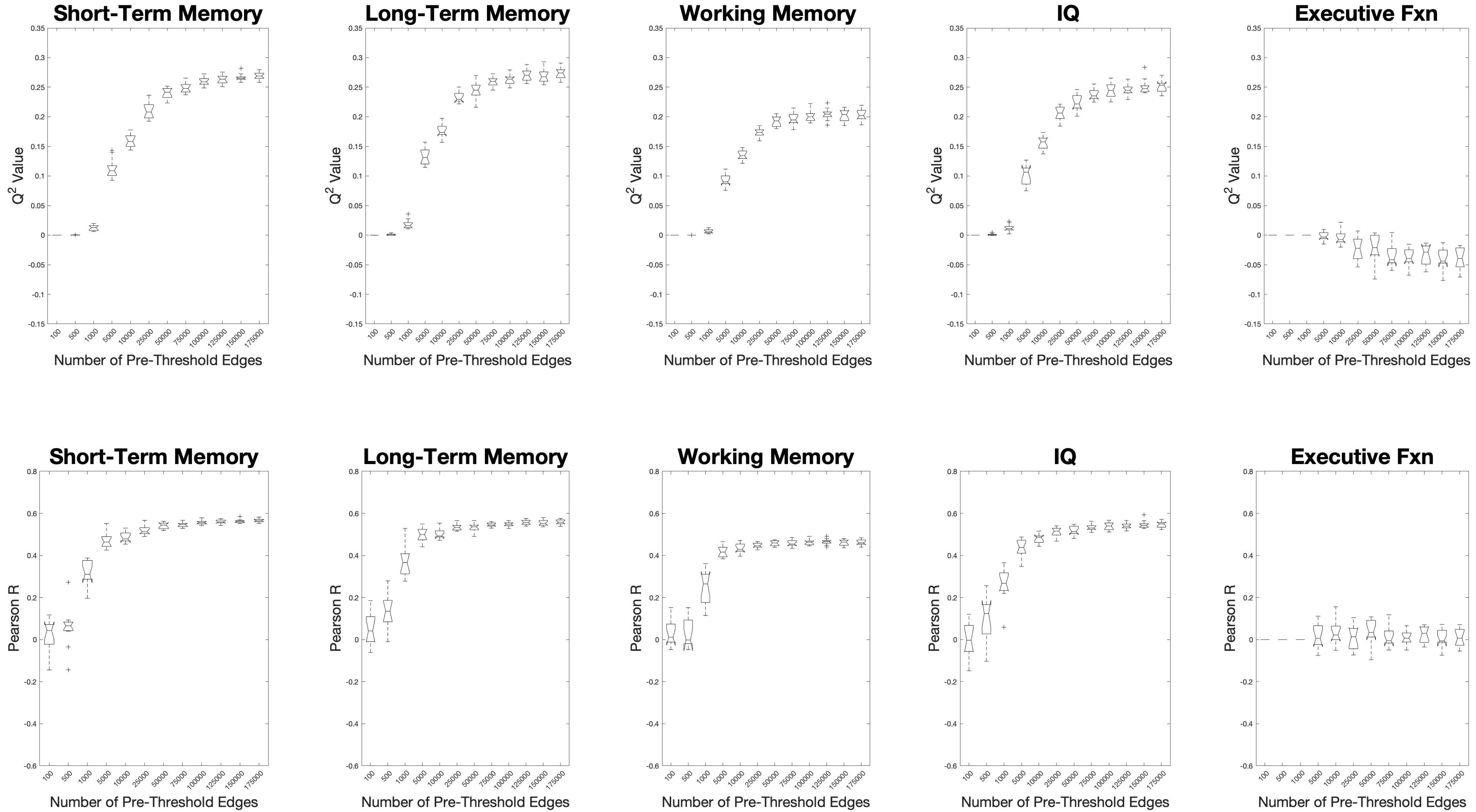

**Supplementary Figure 4. Number of edges as a function of edge threshold.** The effect of different thresholds on the number of significant edges used in ridge regression analyses. As expected, as the threshold becomes more conservative, fewer edges are used in final predictive analysis. See Figures 2 and 3 (main text) and Supplementary Figures 4-5 for how the number of edges and the statistical threshold affects model performance.

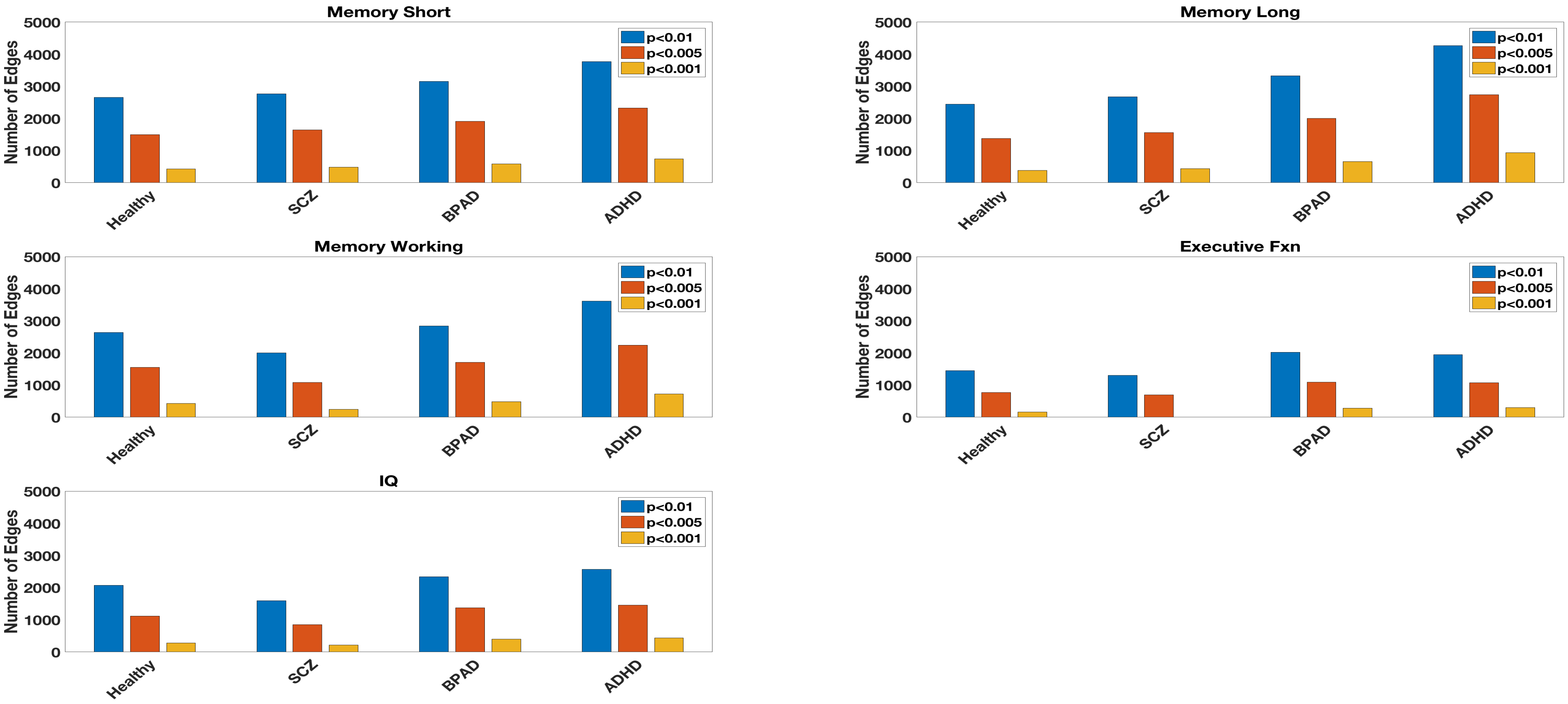

**Supplementary Figure 5.** Performance as a function of edge threshold for 10-fold ridge regression analysis. Each increment select the most significant (smallest p-value) 5, 10, etc. edges. Top row shows performance measured with Q-Squared. Bottom Row shows performance measured as Pearson correlation between actual and predicted cognitive construct scores. We iterated analyses 200 times to produce error bars; we did not regress out motion in these analyses.

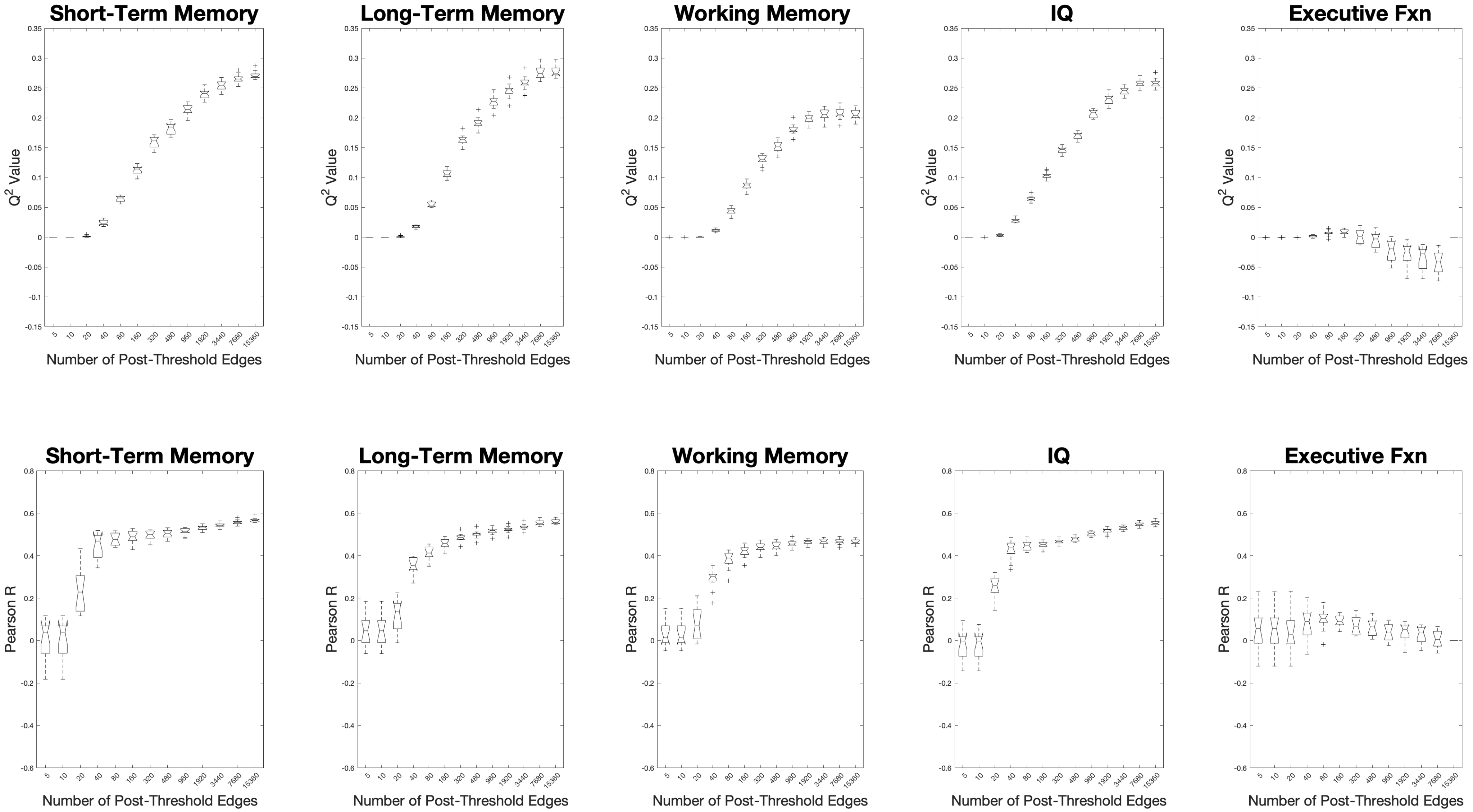

**Supplementary Figure 6.** The effect of motion on model performance. We ran the k-10 fold analyses 1,000 times with and without partial correlation for motion. Except for increased performance on IQ prediction with partial correlation, there was no notable difference in performance with the partial correlation analysis with motion.

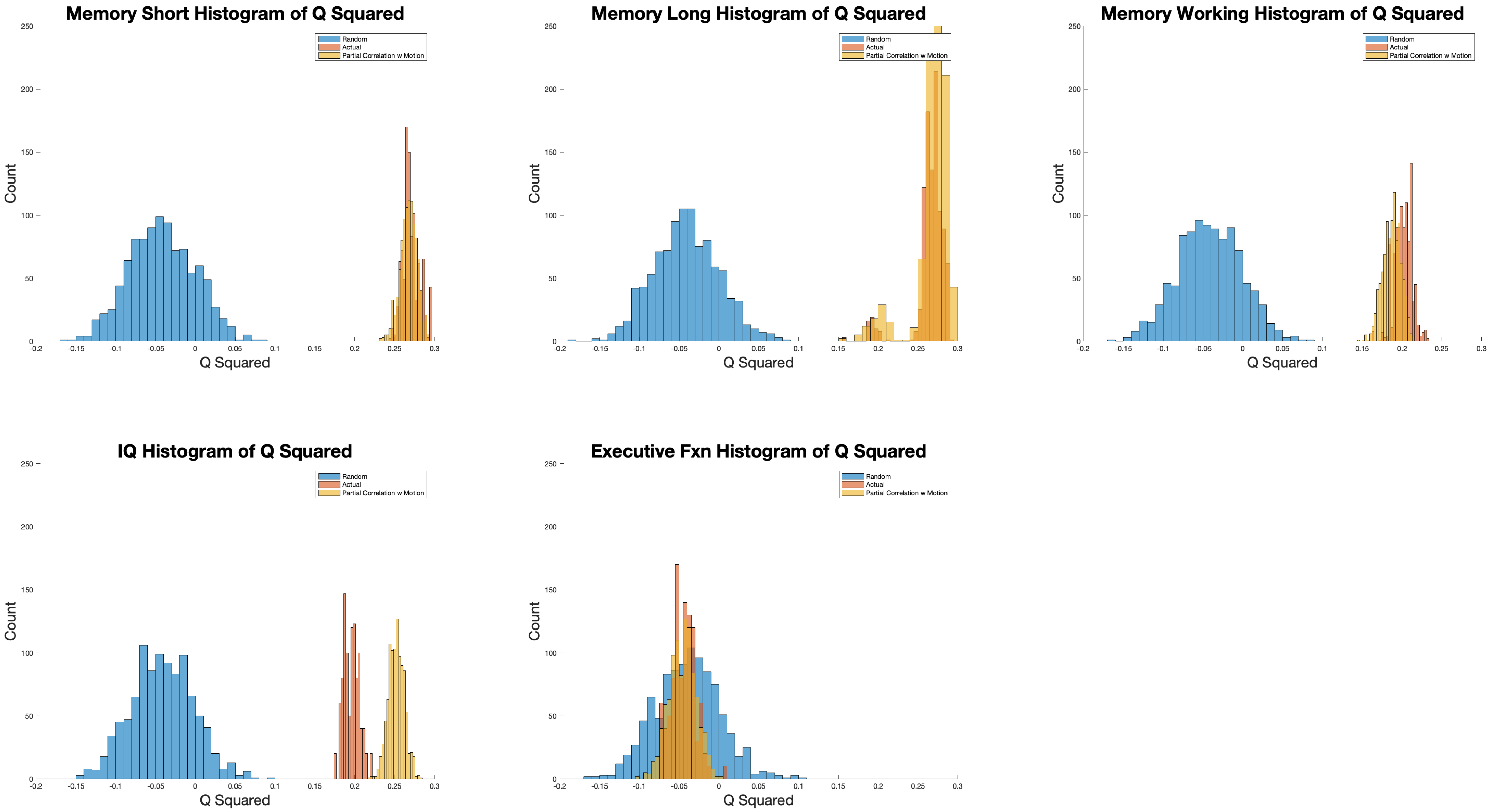

**Supplementary Figure 7. Breakdown of weighted edges by task.** For each task, this figure shows how many edges were weighted by the ridge regression algorithm in all 10 folds of 75% of the 1,000 iterations.

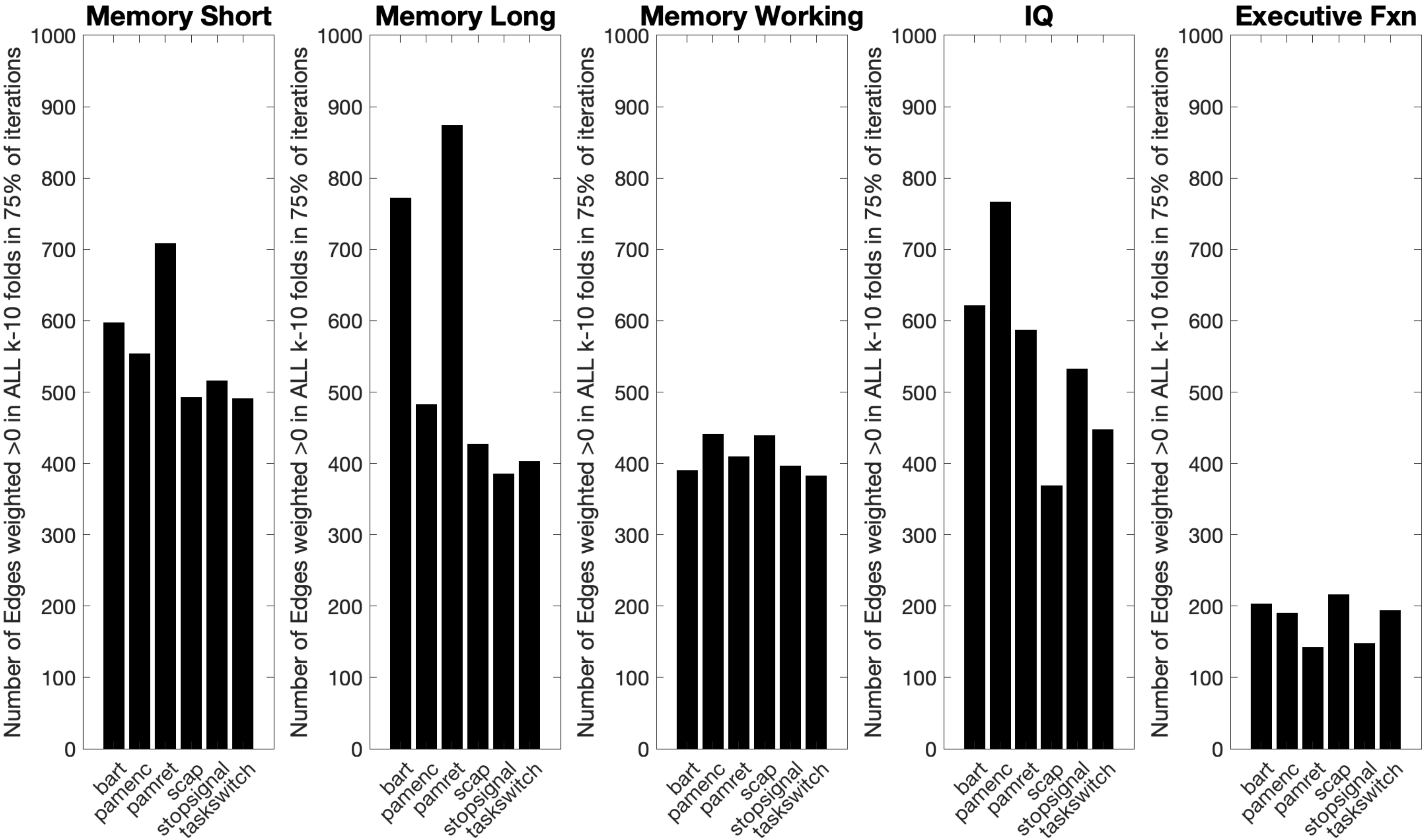

**Supplementary Figure 8. Leave-group-out model performance as a function of edge threshold**. Measures for the leave-one-group-out analysis at p<0.01 (above, right), p<0.005 (below, left) and p<0.001 (below, right) thresholds. As expected, more rigorous thresholds decrease the number of edges and, therefore model performance; however the general trend remains unchanged.

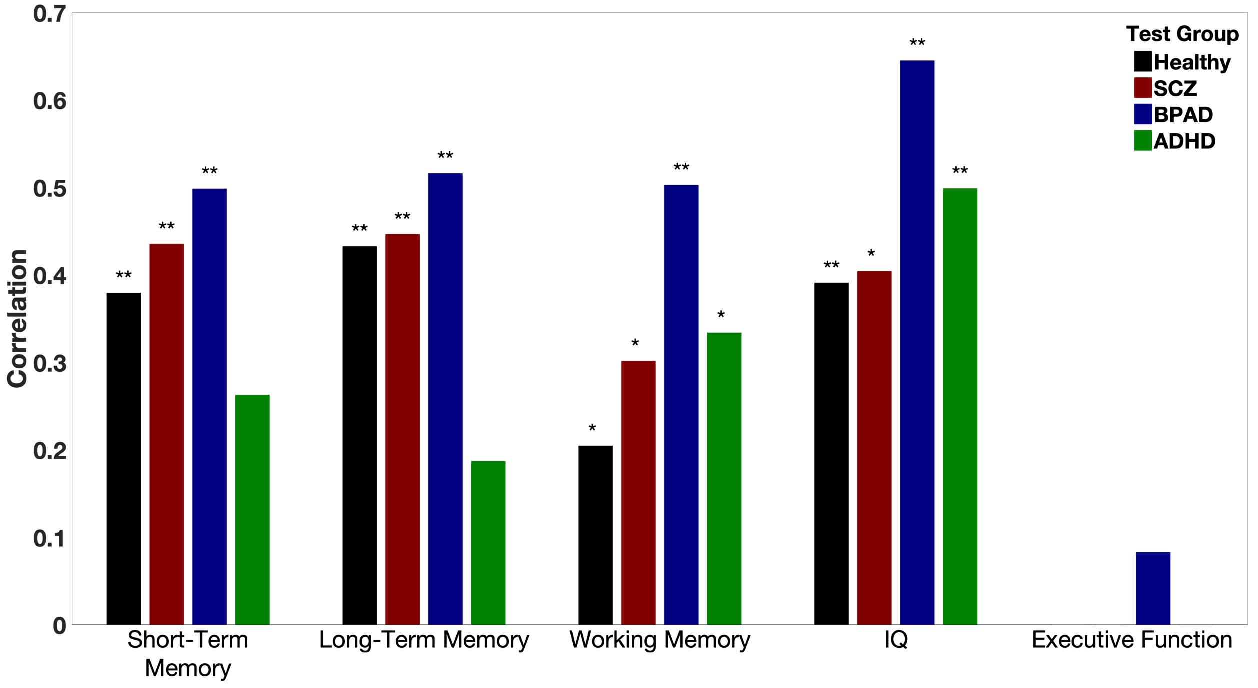


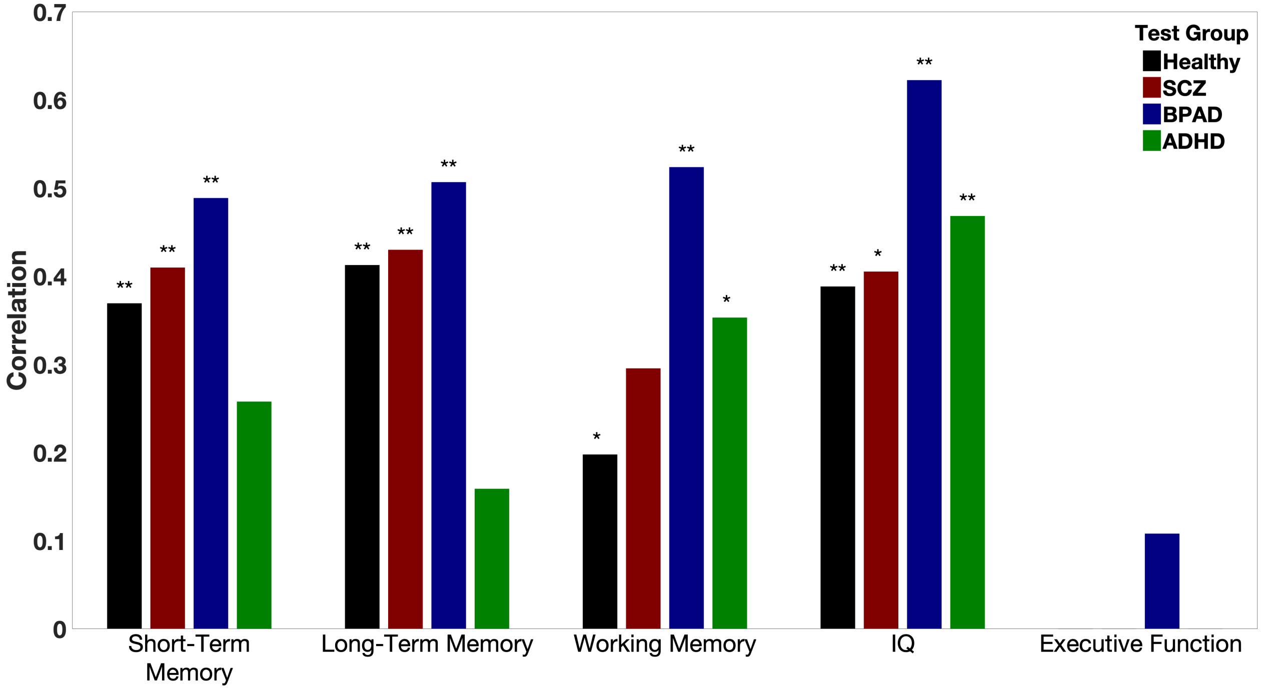


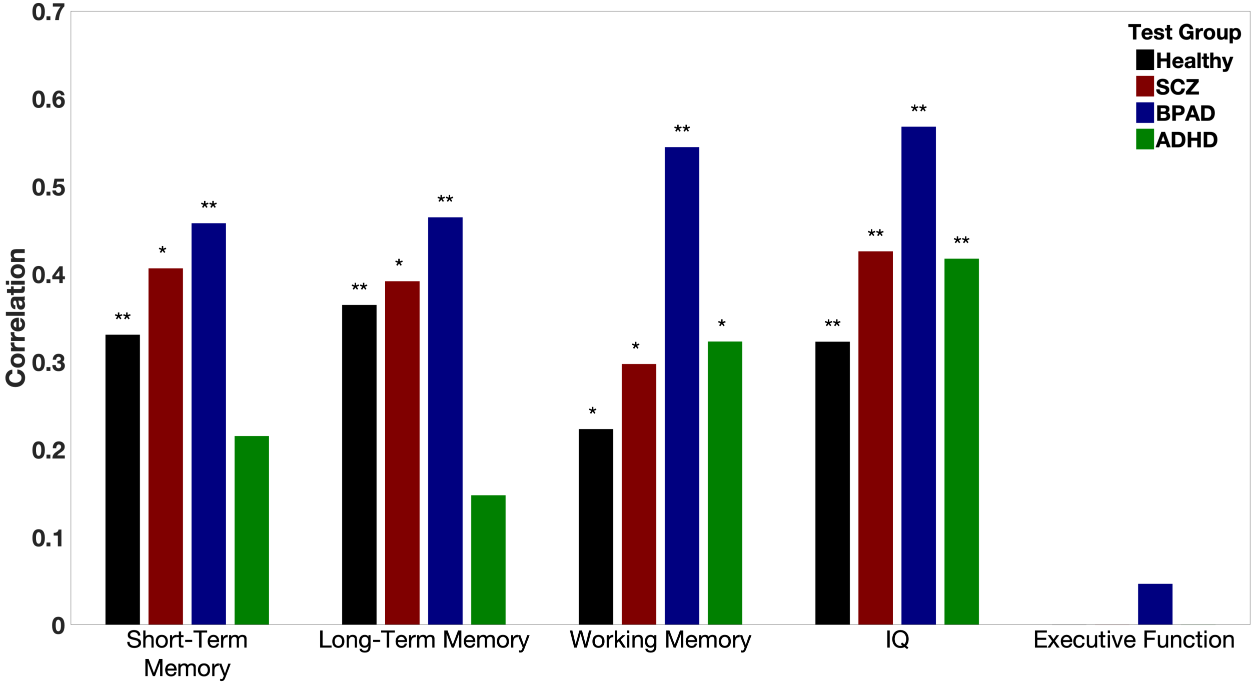

**Supplementary Figure 9A. Short-Term Memory Phenotype Characteristics.** The top row shows boxplots of the individual behavioral measures (as titled) organized by clinical group. These behavioral measures were normalized and combined through principal components analysis. The bottom row shows the percent of variance explained by each of the principal component scores (left) and a boxplot of the first principal component score for each clinical group (right). CVLT=California Verbal Learning Task.

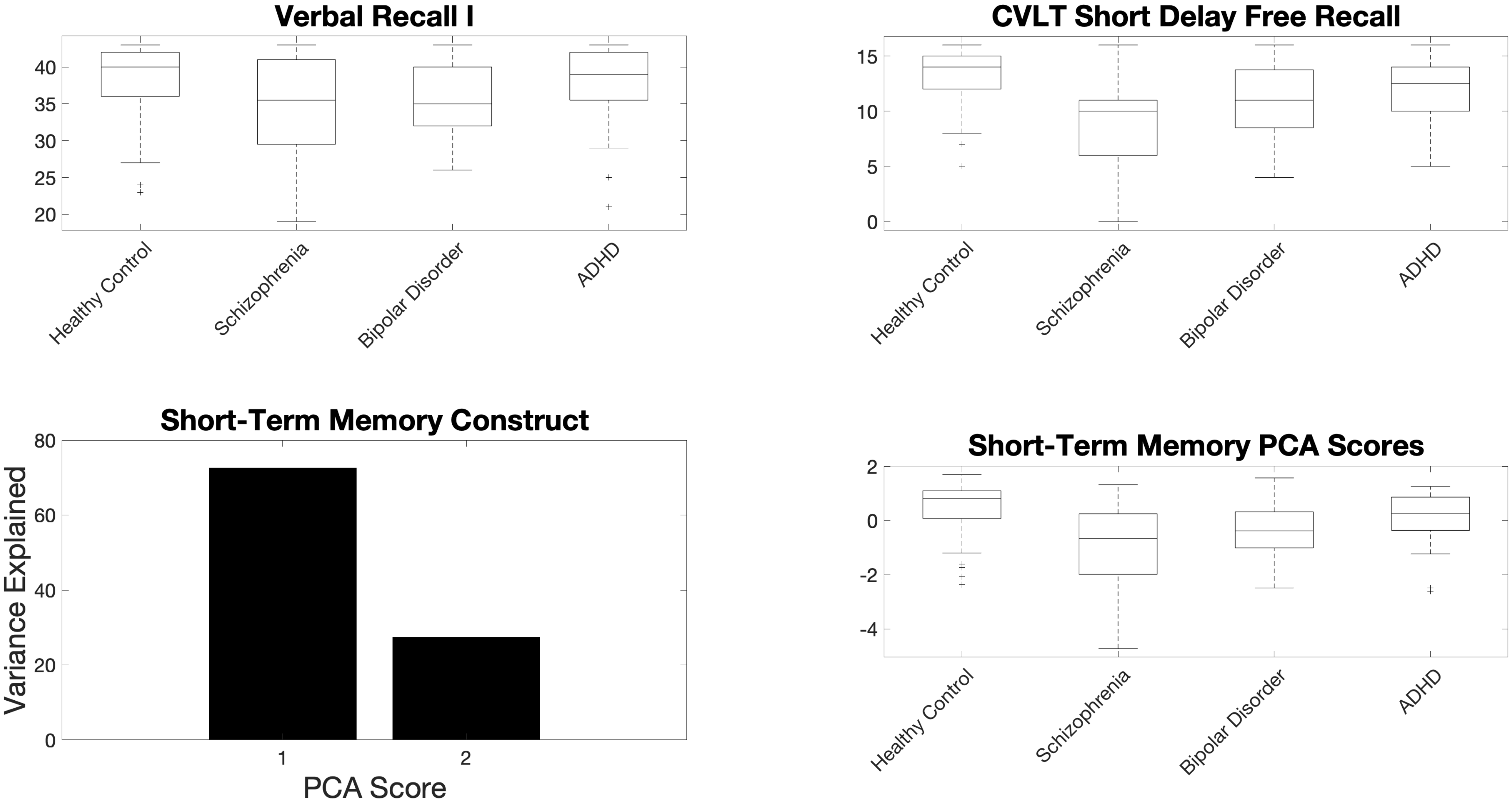

**Supplementary Figure 9B. Long-Term Memory Phenotype Characteristics.** The top row shows boxplots of the individual behavioral measures (as titled) organized by clinical group. These behavioral measures were normalized and combined through principal components analysis. The bottom row shows the percent of variance explained by each of the principal component scores (left) and a boxplot of the first principal component score for each clinical group (right). CVLT=California Verbal Learning Task.

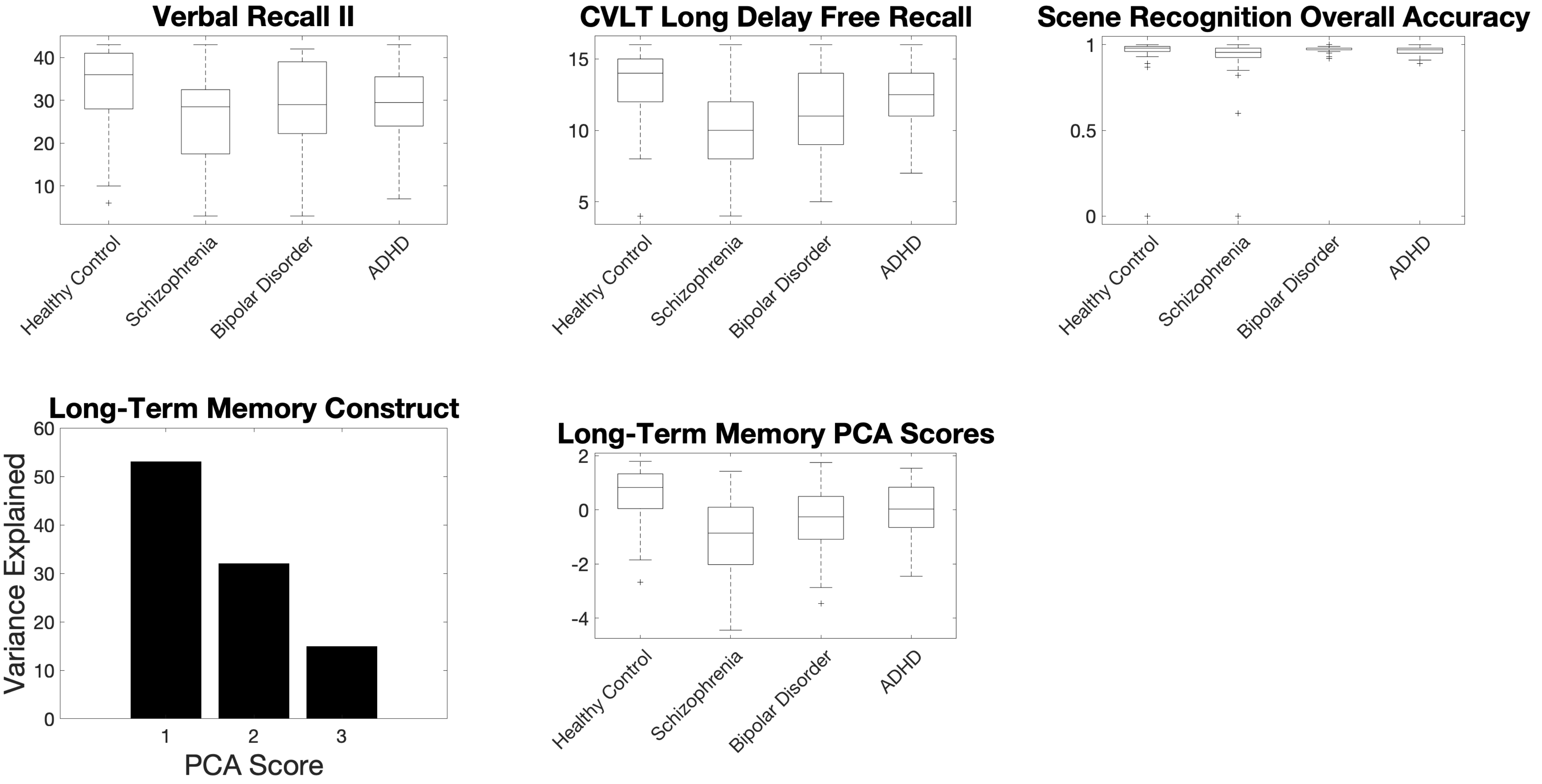

**Supplementary Figure 9C. Working Memory Phenotype Characteristics.** The top row shows boxplots of the individual behavioral measures (as titled) organized by clinical group. These behavioral measures were normalized and combined through principal components analysis. The bottom row shows the percent of variance explained by each of the principal component scores (left) and a boxplot of the first principal component score for each clinical group (right). WMS=Weschler Memory Scale; WAIS=Weschler Adult Intelligence Scale.

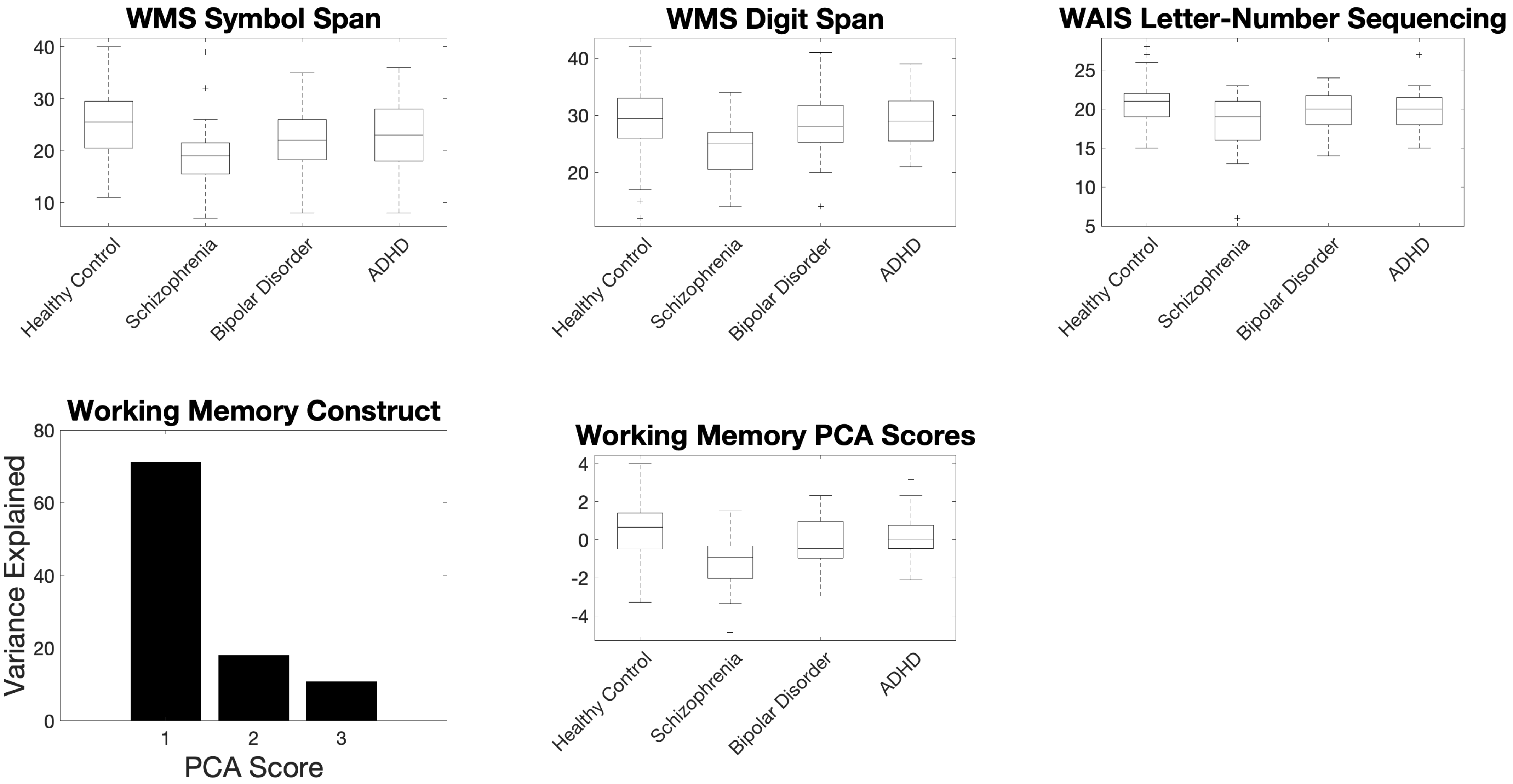

**Supplementary Figure 9D. IQ Phenotype Characteristics.** The top row shows boxplots of the individual behavioral measures (as titled) organized by clinical group. These behavioral measures were normalized and combined through principal components analysis. The bottom row shows the percent of variance explained by each of the principal component scores (left) and a boxplot of the first principal component score for each clinical group (right). WMS=Weschler Memory Scale; WAIS=Weschler Adult Intelligence Scale; IQ=Intelligence Quotient.

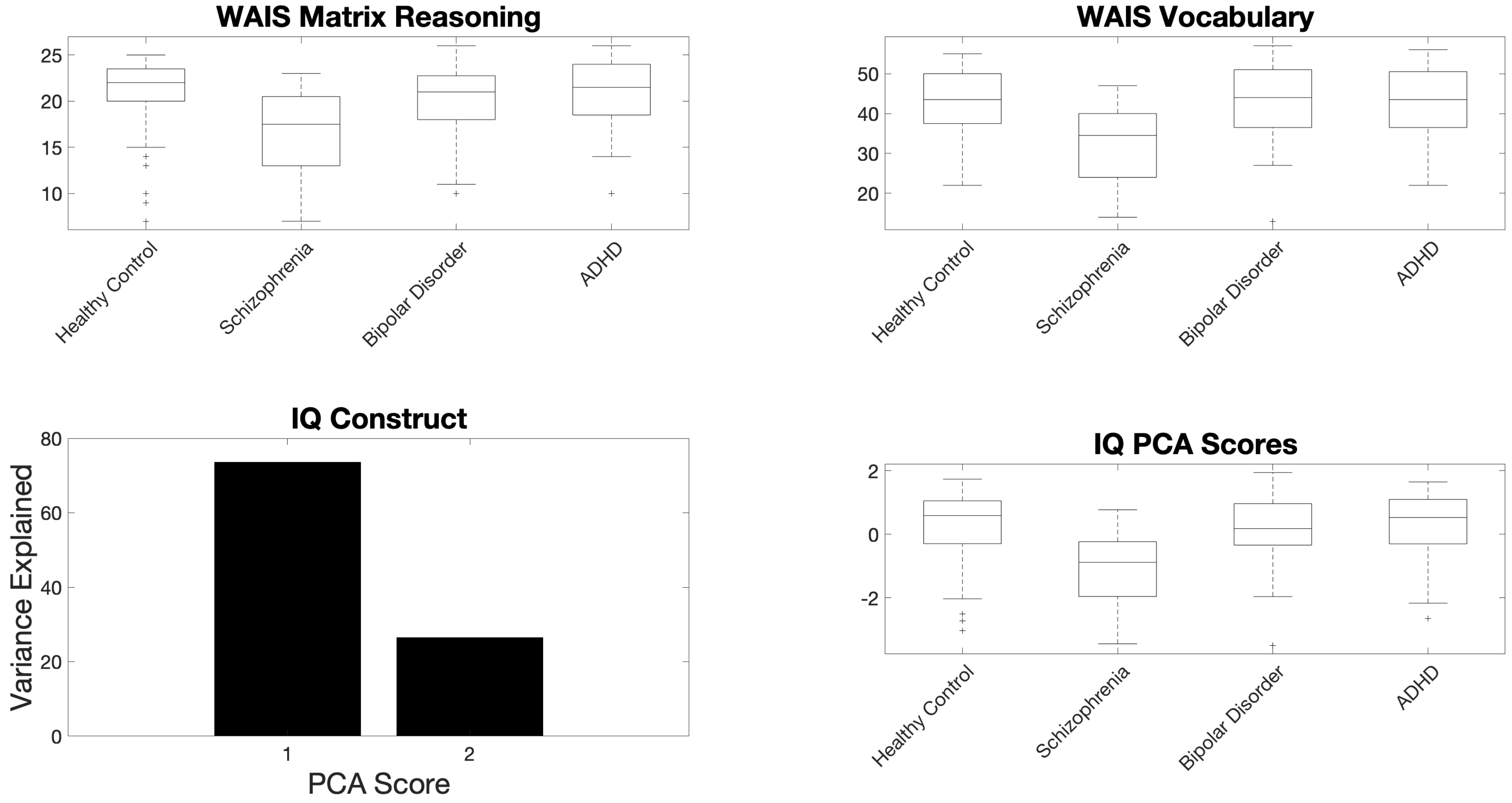

**Supplementary Figure 9E. Executive Function Phenotype Characteristics.** The top row shows boxplots of the individual behavioral measures (as titled) organized by clinical group. These behavioral measures were normalized and combined through principal components analysis. The bottom row shows the percent of variance explained by each of the principal component scores (left) and a boxplot of the first principal component score for each clinical group (right).

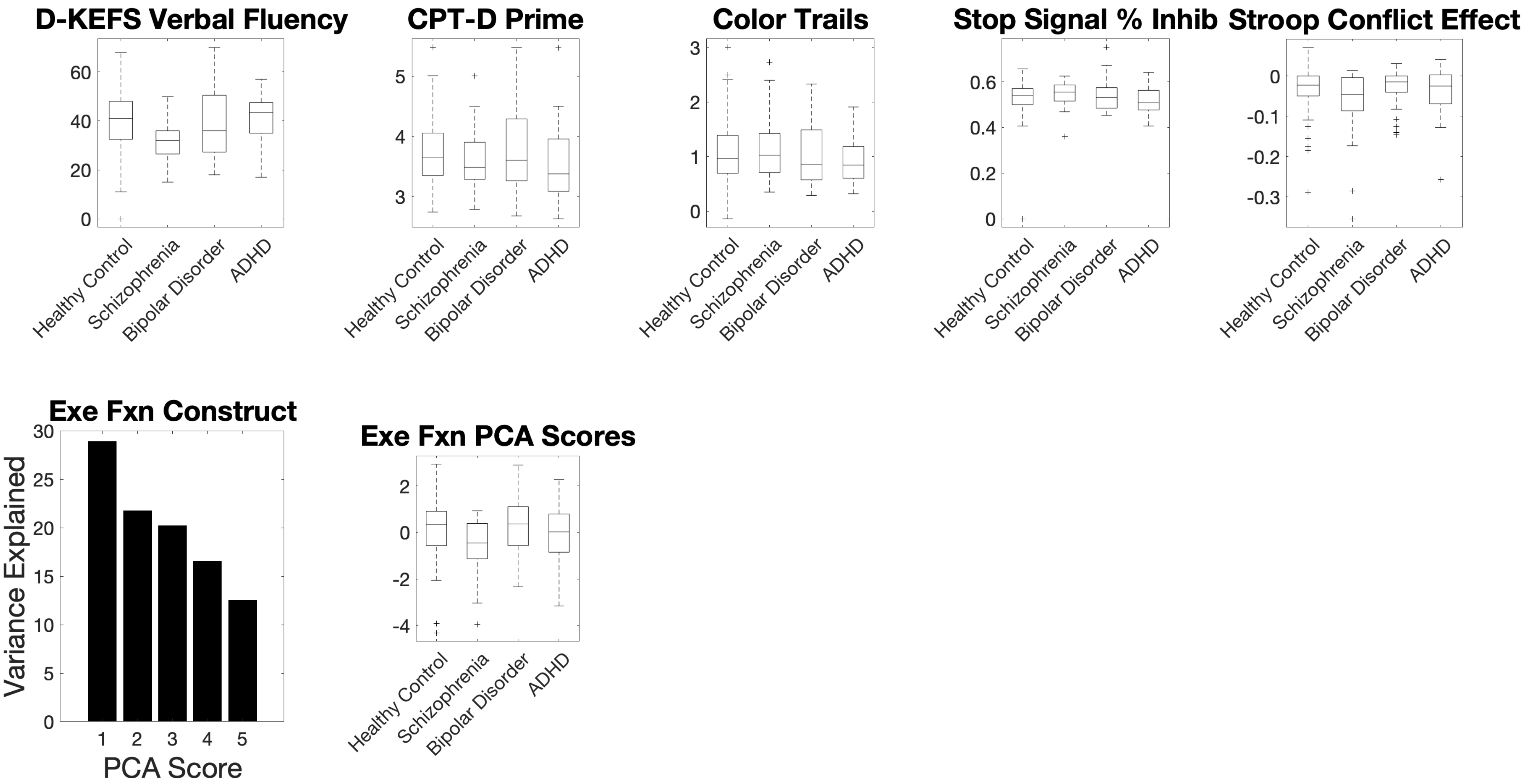

**Supplementary Figure 10. Participant demographics.** Left: brief summary of participant demographics. Cognitive testing scores can be visualized in Figure 2. Right: Venn diagram of medication class. Antipsychotics are represented in red; antidepressants in green, mood stabilizers in blue.

| | Original Number | Excluded | Included Number | Men | Women | Age +/- Stdev | Years Education | Prescribed Antipsychotics | Prescribed Antidepressants | Prescribed Mood Stabilizers |
|---|---|---|---|---|---|---|---|---|---|---|
| Controls | 130 | 54 | 76 | 41 | 35 | 30 +/-8 | 15 +/-1 | 0 | 0 | 0 |
| SCZ | 50 | 18 | 32 | 24 | 8 | 34 +/-9 | 13 +/-1 | 27 | 10 | 7 |
| BPAD | 49 | 14 | 35 | 17 | 18 | 35 +/-9 | 15 +/-2 | 18 | 11 | 23 |
| ADHD | 43 | 11 | 32 | 17 | 15 | 31 +/-10 | 15 +/-2 | 1 | 3 | 1 |
| TOTAL | 272 | 97 | 175 | 99 | 76 | | | 46 | 24 | 31 |

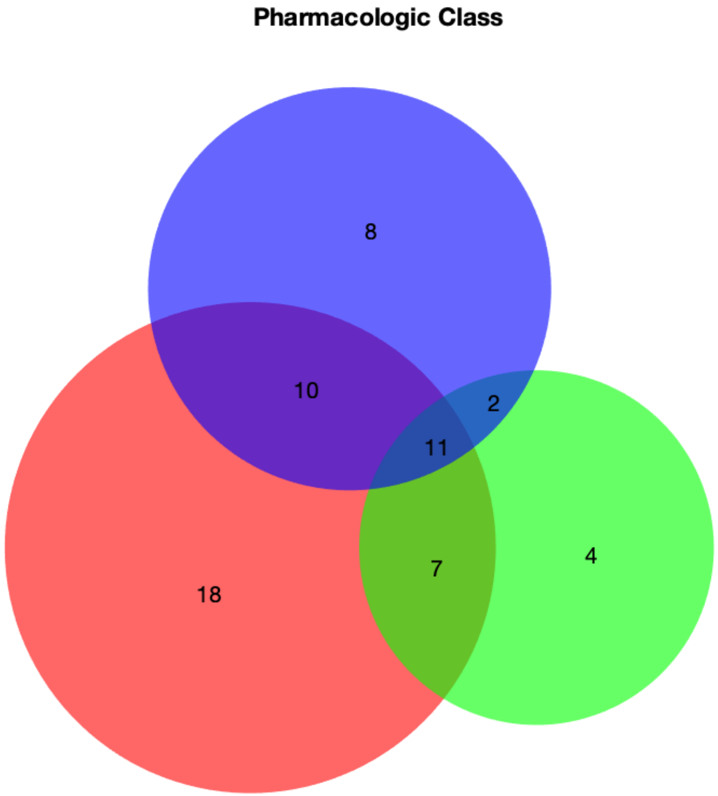

**Supplementary Figure 11.A.** Circle plots showing significantly associated edges for each disease group, broken down by task. Results across all tasks are show in Figure 4 (Main Text).

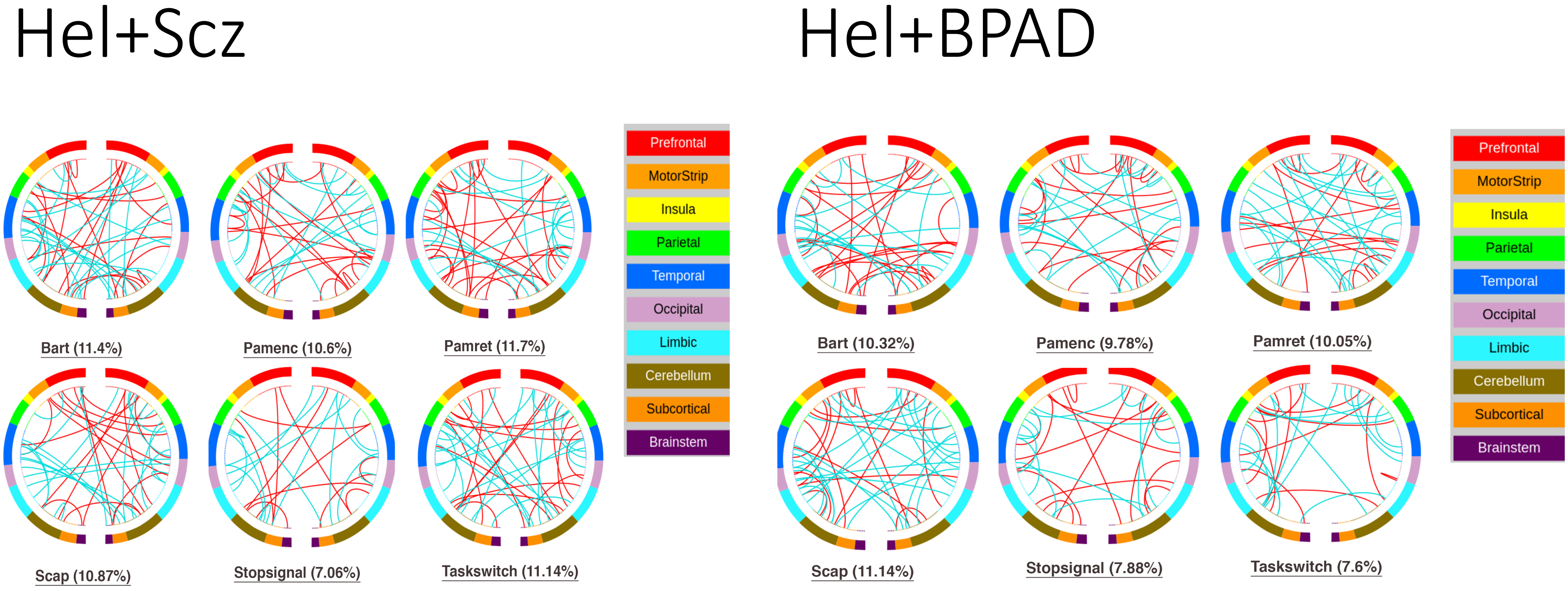

**Supplementary Figure 11.B.** Circle plots showing significantly associated edges for each disease group, broken down by task. Results across all tasks are show in Figure 4 (Main Text).

### Hel+ADHD

### Scz+BPAD

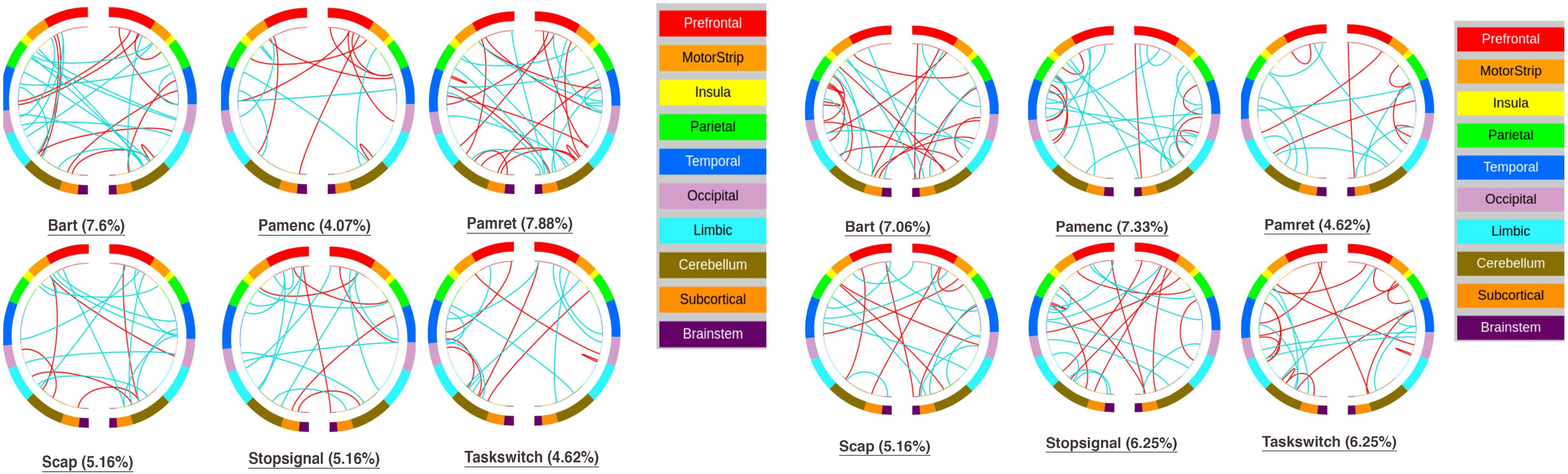

**Supplementary Figure 11.C.** Circle plots showing significantly associated edges for each disease group, broken down by task. Results across all tasks are show in Figure 4 (Main Text).

### Scz+ADHD

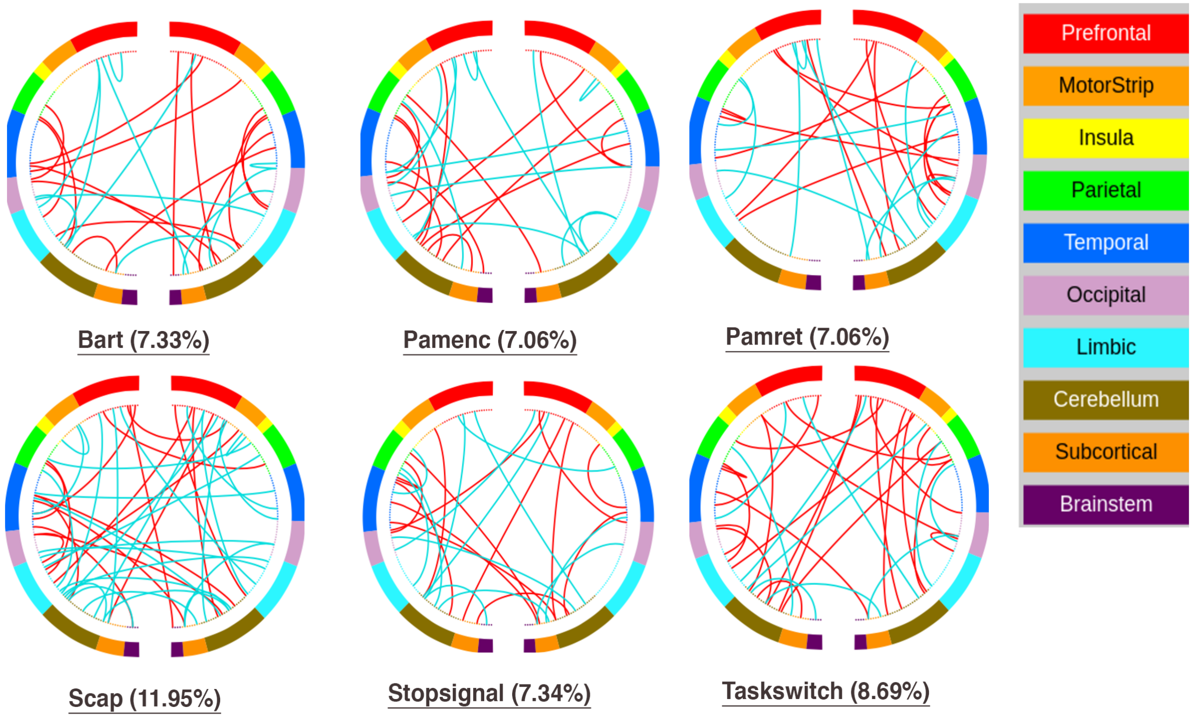


### BPAD+ADHD

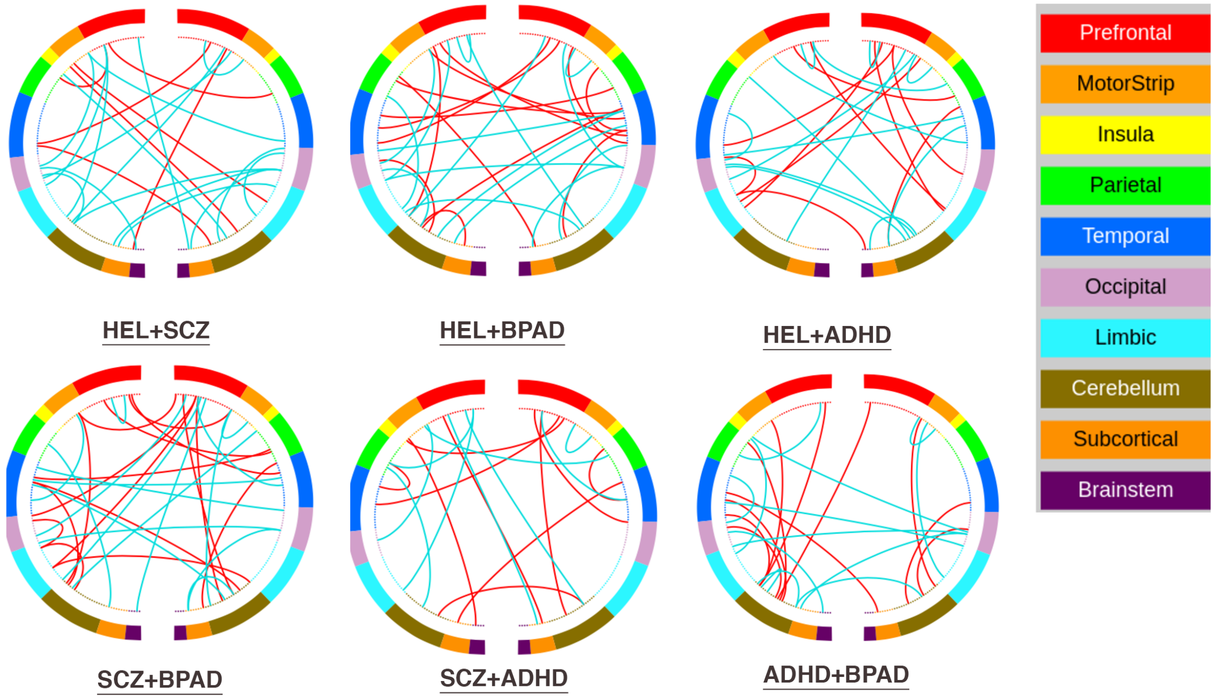
